# A novel SUN1-ALLAN complex coordinates segregation of the bipartite MTOC across the nuclear envelope during rapid closed mitosis in Plasmodium

**DOI:** 10.1101/2024.12.04.625416

**Authors:** Mohammad Zeeshan, Igor Blatov, Ryuji Yanase, David J. P. Ferguson, Sarah L. Pashley, Zeinab Chahine, Yoshiki Yamaryo Botté, Akancha Mishra, Baptiste Marché, Suhani Bhanvadia, Molly Hair, Sagar Batra, Robert Markus, Declan Brady, Andrew Bottrill, Sue Vaughan, Cyrille Y. Botté, Karine Le Roch, Anthony A. Holder, Eelco C. Tromer, Rita Tewari

**Author notes:** These authors contributed equally (Mohammad Zeeshan, Igor Blatov). These authors contributed equally (Ryuji Yanase, David J. P. Ferguson).

## Abstract

Mitosis in eukaryotes involves reorganization of the nuclear envelope (NE) and microtubule-organizing centres (MTOCs). During male gametogenesis in Plasmodium, the causative agent of malaria, mitosis is exceptionally rapid and highly divergent. Within 8 min, the haploid male gametocyte genome undergoes three replication cycles (1N to 8N), while maintaining an intact NE. Axonemes assemble in the cytoplasm and connect to a bipartite MTOC-containing nuclear pole (NP) and cytoplasmic basal body, producing eight flagellated gametes. The mechanisms coordinating NE remodelling, MTOC dynamics, and flagellum assembly remain poorly understood.

We identify the SUN1-ALLAN complex as a novel mediator of NE remodelling and bipartite MTOC coordination during *Plasmodium* male gametogenesis. SUN1, a conserved NE protein, localizes to dynamic loops and focal points at the nucleoplasmic face of the spindle poles. ALLAN, a divergent allantoicase, has a location like that of SUN1, and these proteins form a unique complex, detected by live-cell imaging, ultrastructural expansion microscopy, and interactomics. Deletion of either SUN1 or ALLAN genes disrupts nuclear MTOC organization, leading to basal body mis-segregation, defective spindle assembly, and impaired spindle microtubule-kinetochore attachment, but axoneme formation remains intact. Ultrastructural analysis revealed nuclear and cytoplasmic MTOC miscoordination, producing aberrant flagellated gametes lacking nuclear material. These defects block development in the mosquito and parasite transmission, highlighting the essential functions of this complex.

## Introduction

Mitosis, the process of eukaryotic cell division, requires the accurate segregation of nuclear and cytoplasmic materials, coordinated by dynamic changes in the nuclear envelope (NE) and microtubule-organizing centres (MTOCs) (Liu and Pellman, 2020). The NE, a double-membrane structure with embedded nuclear pore (NP) complexes, is a selective barrier, facilitating signal exchange between cytoplasm and nucleus. The NE consists of inner (INM) and outer (ONM) membranes, with the ONM continuous with the endoplasmic reticulum (ER). Beyond structural and transport roles, the NE is integral to mitosis, supporting spindle formation, kinetochore attachment, and chromosome segregation, while also accommodating changes in ploidy (Dey and Baum, 2021; Smoyer and Jaspersen, 2014).

In most mammalian cells, mitosis is considered ‘open’, characterized by complete NE disassembly, chromosome condensation and segregation via a microtubule-based bipolar spindle. In contrast, in unicellular organisms such as the budding yeast *Saccharomyces cerevisiae,* mitosis is comparatively ‘closed’: the NE remains intact, and chromosomes are segregated by an intranuclear spindle anchored to acentriolar NPs (Boettcher and Barral, 2013; Sazer et al., 2014). However, it is now believed that no mitosis is wholly ‘open’ or ‘closed’ and that in most cases remnants of the NE interact with the spindle to support chromosome segregation (Dey and Baum, 2021). Closed mitosis involves diverse NE dynamics: for example, *Trypanosoma* and *Saccharomyces* employ intranuclear spindle assembly, whereas *Chlamydomonas* and *Giardia* assemble spindles outside the nucleus; the NE remains largely unbroken, but large polar fenestrae are formed to allow microtubule access to the chromosomes (Makarova and Oliferenko, 2016). These examples highlight the fact that ‘open’ and ‘closed’ mitosis are merely two extremes of a continuum of NE remodelling strategies during mitosis, underlining extensive diversity in cell division, a core cellular process.

The interaction between nucleus, the chromatin and in general the nucleolplasmic environment and the cytoskeleton is facilitated by the LINC (Linker of Nucleoskeleton and Cytoskeleton) complex (**Fig. 1**), which bridges the INM and ONM in most eukaryotes via interactions between SUN (Sad1, UNC84)-domain proteins on the INM and KASH (Klarsicht, ANC-1, Syne Homology)-domain proteins on the ONM (Hao and Starr, 2019; Starr and Fridolfsson, 2010). The LINC complex links chromatin and the cytoskeleton throughout eukaryotes (**Fig. 1**). For instance, LINC complexes anchor telomeres to the NE during meiosis in budding yeast (Schober et al., 2008) and cluster centromeres near the INM in fission yeast (Funabiki et al., 1993). Furthermore, SUN proteins connect to the lamin-like proteins that cover the INM on the nucleoplasmic side in animals and plants (Koreny and Field, 2016). In plants, SUN–WIP complexes (analogous to the SUN-KASH proteins of animals and fungi) connect the nucleus to actin filaments (Zhou et al., 2015), while Dictyostelium uses SUN1 to anchor the spindle pole body-like MTOC at the NE (Xiong et al., 2008). Such interactions coordinate nuclear positioning, chromosome organization, and mechanical integration with the cytoskeleton. Evolutionary studies suggest that LINC was established before the emergence of the last eukaryotic common ancestor (LECA), highlighting its ancient role in NE formation and cytoskeletal coordination (**Fig. 1**) (Baum and Baum, 2014; Koreny and Field, 2016). However, as SUN proteins are broadly conserved in many eukaryotic lineages, KASH and lamin-like proteins are often not detected (Koreny and Field, 2016). Intriguingly, in many non-model eukaryotes like for instance apicomplexan parasites (e.g. plasmodium and toxoplasma) microscopic observations reveal chromatin structures such as centromeres, telomeres and the spindle pole body-like MTOCs are to be closely associated with and/or embedded in the NE (**Fig. 1**). *Have KASH/lamins proteins been lost or have novel systems evolved to support a SUN protein-based functions at the nuclear envelope?*

**Fig 1.**
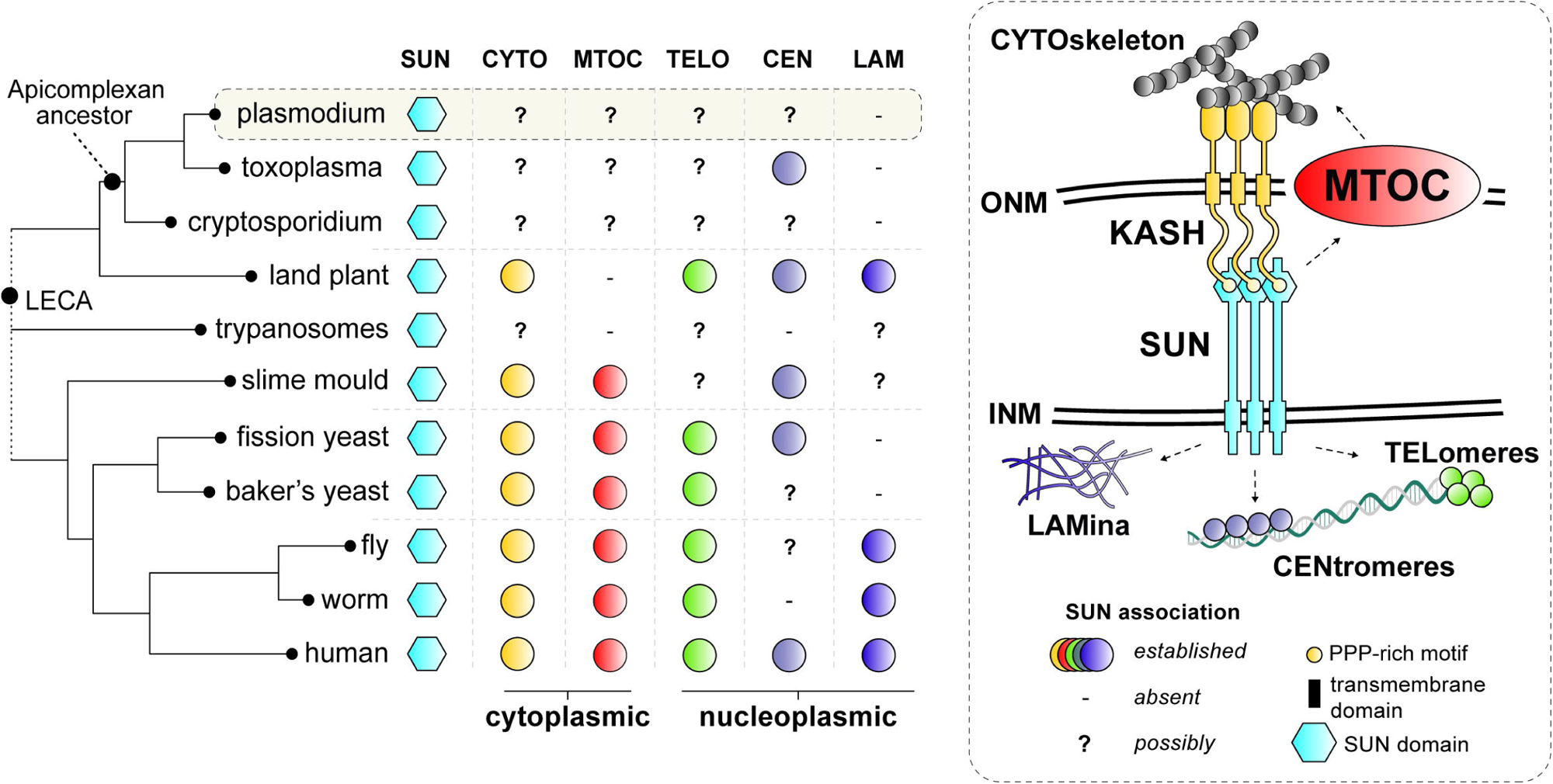
Comparative analysis of SUN protein functions in common eukaryotic model systems. SUN proteins bridge the outer (ONM) and inner (INM) membranes of the nuclear envelope (NE) to link the cytoskeleton (i.e. actin, microtubules and its organising centres) to various heterochromatic domains (i.e. centromere, telomeres) and the nuclear lamina (five different functions with a unique colour). Established roles of SUN proteins in common model organisms are depicted by coloured circles. ‘-‘ no functional connection for SUN was found and/or structures are not present. ‘?’ denotes possible roles for SUN proteins as NE connections with such structures have been established in these lineages. LECA: Last Eukayotic Common Ancestor.

The malaria-causing parasite *Plasmodium spp.* uses closed mitosis with some highly divergent features across its complex life cycle. In the asexual stage in the blood of its vertebrate host, during schizogony there is asynchronous nuclear division without coincident cytokinesis, and during male gametogenesis in the mosquito host there is a unique and rapid form of closed mitosis essential for parasite transmission (Guttery et al., 2022; Sinden et al., 1978). Following activation in the mosquito gut, the haploid male gametocyte undergoes three rounds of genome replication (from 1N to 8N) within just 6 to 8 min, while maintaining an intact NE. Concurrently, axonemes form in the cytoplasm, emanating from bipartite MTOCs that consist of intranuclear spindle poles and cytoplasmic basal bodies (BB). This rapid karyokinesis and subsequent cytokinesis results in the production of eight flagellated haploid male gametes within 15 min (Guttery *et al*., 2022).

The speed of male gametogenesis in *Plasmodium* imposes unique requirements and constraints on cellular structures. Notably, the flagella lack intraflagellar transport (IFT), which is atypical (Sinden et al., 2010). The bipartite organization of the MTOC was recently revealed, and use of fluorescently tagged markers such as SAS4 and kinesin-8B has illuminated the dynamics of BB and axoneme formation (Zeeshan et al., 2022a; Zeeshan et al., 2019a). Kinetochore proteins like NDC80 display an unconventional, largely clustered linear organization on the spindle, redistributing only during successive spindle duplication (Zeeshan et al., 2020). Remarkably, successive spindle and BB segregation occurs within 8 min of gametocyte activation, without NE breakdown, indicating an unusual, streamlined closed mitotic process (Zeeshan *et al*., 2022a; Zeeshan *et al*., 2019a).

The mechanisms that coordinate the formation and function of the intranuclear spindle poles, the cytoplasmic BB, and the NE remain unclear. Specifically, how the NE is remodelled to accommodate rapid genome replication, and how it affects the organization and function of the bipartite MTOC remain open questions. We addressed these questions by investigating NE remodelling and MTOC coordination in *Plasmodium berghei* (Pb), using the conserved NE protein SUN1. By combining live-cell imaging, ultrastructural expansion microscopy (U-ExM), and proteomic analysis, we identify SUN1 as a key NE component. Using different mitotic and MTOC/BB markers, we investigate their coordination and the flexibility of NE during rapid mitosis. Our findings reveal that SUN1 interacts with a novel allantoicase-like protein (termed ALLAN), to form a divergent LINC-like complex without KASH proteins. Functional disruption of either SUN1 or ALLAN impairs BB segregation, disrupts spindle-kinetochore attachment, and results in defective flagellum assembly. Our results highlight a unique divergence of NE remodelling and MTOC organization in *Plasmodium* gametogenesis from conventional eukaryotic models system found in animals, fungi and plants.

## Results

### Generation and Validation of PbSUN1 Transgenic Lines

To investigate the role of PbSUN1 (PBANKA_1430900) during *P. berghei* male gametogenesis, we generated transgenic parasite lines expressing a C-terminal GFP-tagged SUN1 protein (SUN1-GFP). The GFP-tagging construct was integrated into the 3′ end of the endogenous *sun1* locus via single-crossover recombination (**Fig. S1A**), and correct integration was confirmed by PCR analysis using diagnostic primers (**Fig. S1B**). Western blot analysis detected SUN1-GFP at the expected size (∼130 kDa) in gametocyte lysates (**Fig. S1C**). The SUN1-GFP line grew normally and progressed through the life cycle, indicating that tagging did not disrupt PbSUN1 function. The SUN1-GFP line was used to study the subcellular localization of PbSUN1 across multiple life cycle stages, and its interaction with other proteins during gametogenesis.

### Spatiotemporal Dynamics of SUN1 During Male Gametogenesis

SUN1 protein expression was undetectable by live-cell imaging during asexual erythrocytic stages of the parasite, but robust expression was visible in both male and female gametocytes following activation in ookinete medium (**Fig. S1D, E**). In activated male gametocytes undergoing mitotic division, SUN1-GFP had a dynamic localization around the nuclear DNA (Hoechst-stained), associated with loops and folds formed in the NE during the transition from 1N to 8N ploidy (**Fig. 2A, Supplementary Video 1**). The NE loops extended beyond the Hoechst-stained DNA, suggesting an abundant non-spherical NE membrane (**Fig. 2A**). To further characterize NE morphology during male gametogenesis, serial block-face scanning electron microscopy (SBF-SEM) was used. Analysis of wild-type male gametocytes revealed a non-spherical, contorted nucleus (**Fig. 2B, S1F**). Thin loops of NE (indicated with arrows) were prominent, consistent with the dynamic and irregular location of SUN1-GFP detected by live-cell imaging. 3D-modelling of gametocyte nuclei further confirmed the irregular, non-spherical structure of the NE (**Fig. 2C, S1G**), which expanded rapidly during the period of genome replication. These findings suggest that the male gametocyte NE is a highly dynamic structure that can expand rapidly to accommodate the increased DNA due to replication during gametogenesis.

**Fig 2.**
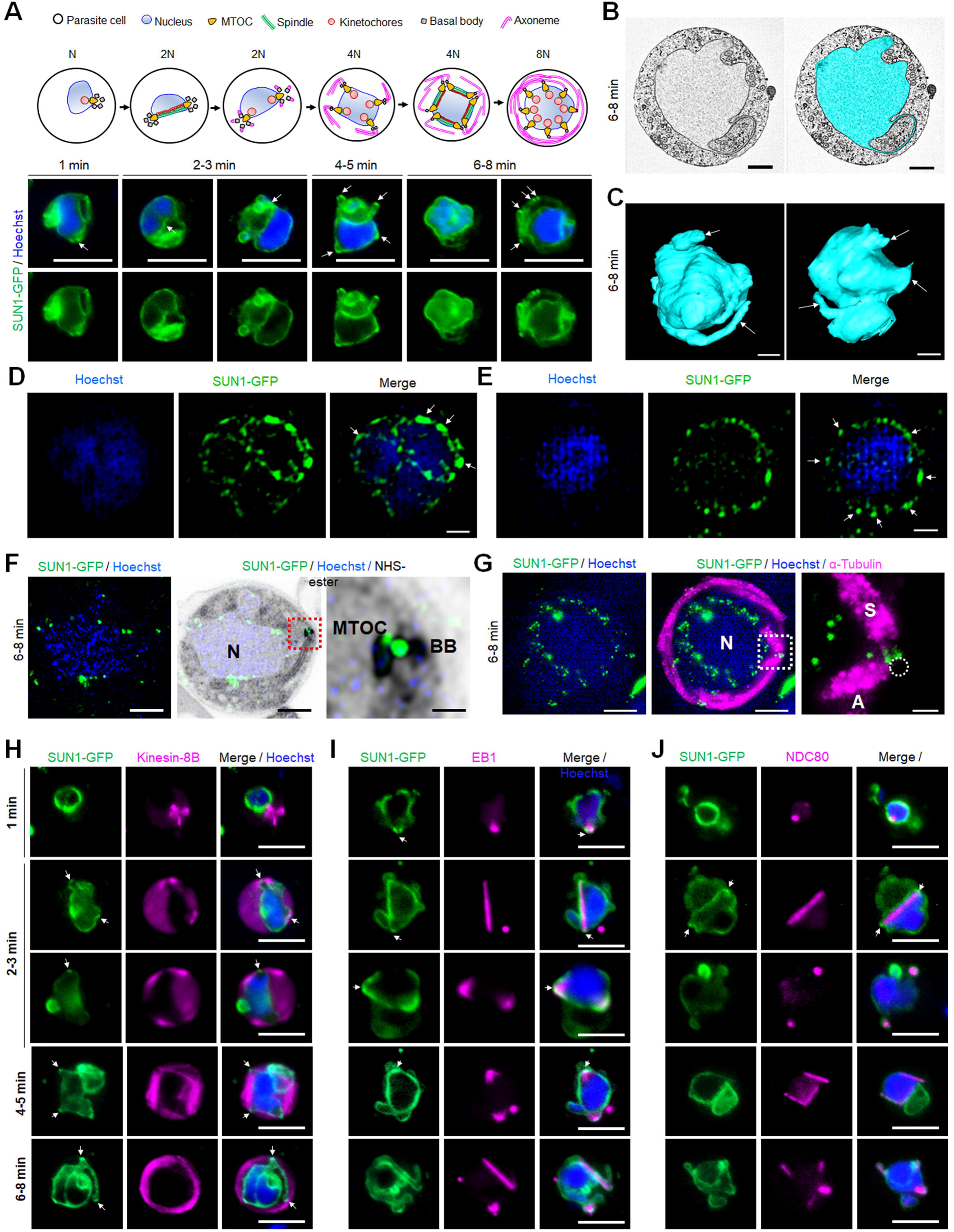
Location of SUN1 during male gametogenesis. **A.** The upper panel schematic illustrates the process of male gametogenesis. N, genome ploidy. Live cell images show the location of SUN1-GFP (green) at different time points(1-8min) during male gametogenesis. DNA(blue) was stained with Hoechst. White arrows indicate the loop/folds. Representative images of more than 50 cells with more than three biological replicates. Scale bar: 5 μm. **B.** Serial block face-scanning electron microscopy (SBF-SEM) data slice of a gametocyte highlighting the complex morphology of the nucleus (cyan). Representative of more than 10 cells. Scale bar: 1 μm. **C.** Two 3D models of gametocyte nuclei showing their contorted and irregular morphology. Representative of more than 10 cells. Scale bar: 1 μm. **D.** SIM images of SUN1-GFP male gametocytes activated for 8 min and fixed with paraformaldehyde. Arrows indicate the SUN1-GFP signals with high intensity after fixation. Representative image of more than 10 cells from more than two biological replicates. Scale: 1 µm. **E.** SIM images of SUN1-GFP male gametocytes activated for 8 min and fixed with methanol. Arrows indicate the SUN1-GFP signals with high intensity after fixation. Representative image of more than 10 cells from more than two biological replicates. Scale: 1 µm. **F.** Expansion microscopy (ExM) images showing location of SUN1 (green) detected with anti-GFP antibody and BB/MTOC stained with NHS ester (grey). Hoechst was used to stain DNA. Scale bar: 5 µm. Inset is the area marked with the red box around the BB/MTOC highlighted by NHS-ester staining. Scale bar: 1LJµm. Representative images of more than 10 cells from two biological replicates. **G.** ExM images showing location of SUN1 (green) and α-tubulin (magenta) detected with anti-GFP and anti-tubulin antibodies, respectively. Hoechst was used to stain DNA(blue). N=Nucleus; S=Spindle; A=Axoneme. Scale bar: 5 µm. Inset is the area marked with the white box on Fig. 1E middle panel around the BB/MTOC. Scale bar: 1LJµm. Representative images of more than 10 cells from two biological replicates. **H.** Live cell imaging showing location of SUN1-GFP (green) in relation to the BB and axoneme marker, kinesin-8B-mCherry (magenta) at different time points(1-5min) during gametogenesis. Blue in merged image is DNA stained with Hoechst. Representative images of more than 20 cells from more than three biological replicates. White arrows indicate the loops/folds labelled with SUN1 where BB/axonemes are assembled outside the nuclear membrane. Scale bar: 5LJµm. **I**. Live cell imaging showing location of SUN1-GFP (green) in relation to the spindle marker, EB1-mCherry (magenta) at different time points during gametogenesis. Blue in merged image is DNA stained with Hoechst. White arrows indicate the loops/folds labelled with SUN1. Representative images of more than 20 cells from more than three biological replicates. Scale bar: 5LJµm. **J**. Live cell imaging showing location of SUN1-GFP (green) in relation to the kinetochore marker, NDC80-mCherry (magenta) at different time points during gametogenesis. Blue in merged image is DNA stained with Hoechst. White arrows indicate the loops/folds labelled with SUN1. Representative images of more than 20 cells with more than three biological replicates. Scale bar: 5LJµm.

### SUN1 Dynamics by High-Resolution Imaging

To resolve the SUN1-GFP location with higher precision, structured illumination microscopy (SIM) was performed on SUN1-GFP gametocytes fixed at 6-8 min post activation. Fixation of gametocytes caused SUN1-GFP signal coalescence into regions within the NE (**Fig. S2A**), but an uneven intensity of SUN1-GFP signal around the DNA was observed, with areas of higher intensity (**Fig. 2D, E; S2B**, white arrows) likely representing collapsed forms of the SUN1-GFP-labelled loops observed by live imaging (**Fig. S2A**). To refine the spatial relationship of SUN1 to the BB and the spindle, ultrastructure expansion microscopy (U-ExM) was used. At 6 to 8 min post-activation SUN1-GFP was prominently located around the DNA and at the junction of the nuclear MTOC and the BB (**Fig. 2F; S3A**). Spindle and axoneme staining with α-tubulin antibodies revealed that SUN1-GFP foci were located next to spindle poles (**Fig. 2G; S3B**), suggesting a role in MTOC organization during mitosis. WT-ANKA gametocytes (with no GFP present) did not react with the anti-GFP antibodies (**Fig. S3C**), confirming the specificity of the GFP signal in SUN1-GFP gametocytes.

### The Location of SUN1 Relative to Key Mitotic Markers During Male Gametogenesis

To investigate the association of SUN1 with MTOCs, mitotic spindles, and BB/axonemes during male gametogenesis, its location was compared in real time with that of three markers: the cytoplasmic BB/axonemal protein kinesin-8B, the spindle microtubule-binding protein EB1, and the kinetochore marker NDC80 (Zeeshan *et al*., 2019a; Zeeshan *et al*., 2020; Zeeshan et al., 2023). A parasite line expressing SUN1-GFP (green) was crossed with lines expressing mCherry (red)-tagged kinesin-8B, EB1, or NDC80, and progeny were analyzed by live-cell imaging to determine the proteins’ spatiotemporal relationships.

Within one-min post-activation, kinesin-8B appeared as a tetrad marking the BB in the cytoplasm and close to the nucleus, while SUN1-GFP localized to the nuclear membrane, with no overlap between the signals (**Fig. 2H**). As gametogenesis progressed, nuclear elongation was observed, with SUN1-GFP delineating the nuclear boundary. Concurrently, the BB (the kinesin-8B-marked tetrad) duplicated and separated to opposite poles in the cytoplasm, outside the SUN1-GFP-defined nuclear boundary (**Fig. 2H**). By 2 to 3 min post-activation, axoneme formation commenced at the BB, while SUN1-GFP marked the NE with small puncta located near the nuclear MTOCs situated between the BB tetrads (**Fig. 2H**).

Analysis of gametogenesis with SUN1-GFP and the spindle microtubule marker EB1, revealed a localized EB1-mCherry signal near the DNA, inside the nuclear membrane (marked by SUN1-GFP), within the first min post-activation (**Fig. 2I**). Loops and folds in the nuclear membrane, marked by SUN1-GFP, partially overlapped with the EB1 focal point associated with the spindle pole/MTOC (**Fig. 2I**, average Pearson Corelation Coefficient, R<0.6). At 2 to 3 min post activation, EB1 fluorescence extended to form a bridge-like spindle structure within the nucleus, flanked by two strong focal points that partially overlapped with SUN1-GFP fluorescence (**Fig. 2I**). A similar dynamic location was observed for another spindle protein, ARK2 (**Fig. S4A**).

Kinetochore marker NDC80 showed a similar pattern to that of EB1. Within one-min post-activation, NDC80 was detected as a focal point inside the nuclear membrane, later extending to form a bridge-like structure that split into two halves within 2 to 3 min, and with no overlap with SUN1-GFP (**Fig. 2J**). As with EB1, the nuclear membrane loops formed around NDC80 focal points, maintaining close proximity to the kinetochore bridge (**Fig. 2J**).

Together, these observations suggest that although SUN1-GFP fluorescence is partially colocated with spindle poles revealed by EB1 fluorescence, there is no overlap with kinetochores (NDC80) or BB/axonemes (kinesin-8B).

### SUN1 Location During Female Gametogenesis and Zygote to Ookinete transformation

The location of SUN1-GFP during female gametogenesis was examined using real-time live-cell imaging. Within one-min post-activation, SUN1-GFP was observed to form a half-circle around the nucleus, eventually encompassing the entire nucleus after six to eight min (**Fig. S4B**). In contrast to SUN1-GFP in male gametocytes, SUN1-GFP in female gametocytes was more uniformly distributed around the nucleus without apparent loops or folds in the NE (**Fig. S4B**). This pattern probably reflects the absence of DNA replication and mitosis in female gametogenesis, so that the nucleus remains compact.

During zygote to ookinete differentiation and oocyst development, in zygotes SUN1-GFP had a spherical distribution around Hoechst-stained nuclear DNA at two hours post-fertilization (**Fig. S4C**), but by 24 hours post-fertilization, as the zygote developed into a banana-shaped ookinete, the SUN1-GFP-marked NE became elongated or oval, possibly due to spatial constraints within the cytoplasm (**Fig. S4C**).

### SUN1 is Essential for Basal Body Segregation and Axoneme-Nucleus Coordination During Male Gametogenesis

The role of SUN1 was assessed by deleting its gene using a double crossover homologous recombination strategy in a parasite line constitutively expressing GFP (WT-GFP) (Janse et al., 2006) (**Fig. S5A**). WT-GFP parasites express GFP at all stages of the life cycle, facilitating phenotypic comparisons. Diagnostic PCR confirmed the correct integration of the targeting construct at the *sun1* locus (**Fig. S5B**), and this was verified by qRT-PCR analysis, which showed complete deletion of the *sun1* gene in the resulting transgenic parasite (Δ*sun1*) (**Fig. S5C**).

Phenotypic analysis of the Δ*sun1* line was conducted across the life cycle, in comparison to a WT-GFP control. Two independent knockout clones (Clone-1 and Clone-4) were examined: the clones exhibited similar phenotypes, and one, Clone-4 was used for further experiments. Δ*sun1* parasites produced a comparable number of gametocytes to WT-GFP parasites, but had reduced male gametogenesis, as evidenced by a significant decrease in gamete formation (exflagellation) (**Fig. 3A**). The differentiation of zygotes into ookinetes was also reduced (**Fig. 3B**).

**Fig 3.**
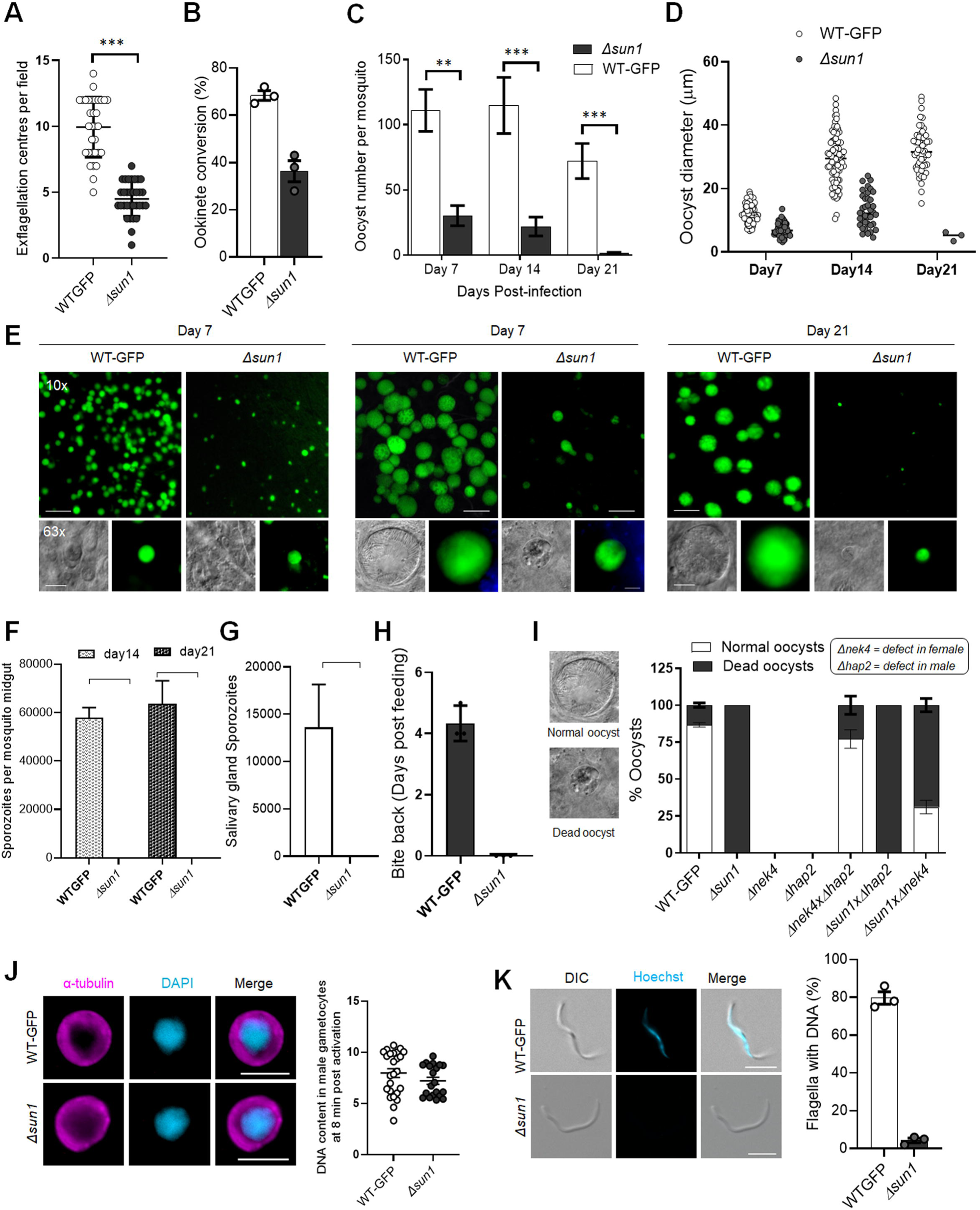
Deletion of *sun1* affects male gamete formation and blocks parasite transmission. **A.** Exflagellation centres per field at 15 min post-activation. n = 3 independent experiments (>10 fields per experiment). Error bar, ± SEM. **B.** Percentage ookinete conversion from zygote. n= 3 independent experiments (> 100 cells). Error bar, ± SEM. **C.** Total number of GFP-positive oocysts per infected mosquito in Δ*sun1* compared to WT-GFP parasites at 7-, 14-and 21-days post infection. Mean ± SEM. n= 3 independent experiments. **D.** The diameter of GFP-positive oocysts in Δ*sun1* compared to WT-GFP parasites at 7-, 14- and 21-days post infection. Mean ± SEM. n= 3 independent experiments. **E.** Mid guts at 10x and 63x magnification showing oocysts of Δ*sun1* and WT-GFP lines at 7-, 14- and 21-days post infection. Scale bar: 50 μm in 10x and 20 μm in 63x. **F.** Total number of midguts sporozoites per infected mosquito in Δ*sun1* compared to WT-GFP parasites at 14- and 21-days post infection. Mean ± SEM. n= 3 independent experiments. **G.** Total number of salivary gland sporozoites per infected mosquito in Δ*sun1* compared to WT-GFP parasites at 21-days post infection. Mean ± SEM. n= 3 independent experiments. **H.** Bite back experiments showing no transmission of Δ*sun1,* while WT-GFP parasites show successful transmission from mosquito to mouse. Mean ± SEM. *n* = 3 independent experiments. **I.** Rescue experiment showing Δ*sun1* phenotype is due to defect in male *sun1* allele. Mean ± SEM. n= 3 independent experiments. **J.** Representative images of male gametocytes at 8 min post activation stained with DAPI and tubulin (left). Fluorometric analyses of DNA content (N) after DAPI nuclear staining (right). The mean DNA content (and SEM) of >30 nuclei per sample are shown. Values are expressed relative to the average fluorescence intensity of 10 haploid ring-stage parasites from the same slide. **K.** Representative images of flagellum (male gamete) stained with Hoechst for DNA (left). The presence or absence of Hoechst fluorescence was scored in at least 30 microgametes per replicate. Mean ± SEM. *n* = 3 independent experiments.

To evaluate sporogonic development, mosquitoes were fed with Δ*sun1* parasites, and oocyst formation was examined. The number of oocysts was markedly reduced in Δ*sun1* parasite-infected mosquitoes compared to WT-GFP parasites at days 7, 14, and 21 post-feeding (**Fig. 3C**). At day 7, oocysts were comparable in size but failed to grow further, and they had degenerated by day 21 (**Fig. 3D, E**) with no evidence of sporozoite production in midgut or salivary glands (**Fig. 3F, G**). Transmission experiments revealed that mosquitoes infected with Δ*sun1* parasites were unable to transmit the parasite to naïve mice; in contrast mosquitoes successfully transmitted the WT-GFP parasite, resulting in a blood-stage infection detected four days later (**Fig 3H**).

Because the defect in Δ*sun1* parasites led to a transmission block, we investigated whether the defect was rescued by restoring *sun1* into Δ*sun1* parasites using either the Δ*nek4* parasite that produces normal male gametocytes but is deficient in production of female gametocytes (Reininger et al., 2005) or the Δ*hap2* parasite that produces normal female gametocytes but is deficient in production of male gametocytes (Liu et al., 2008). We performed a genetic cross between Δ*sun1* parasites and the other mutants deficient in production of either male (Δ*hap2)* or female (Δ*nek4)* gametocytes. Crosses between Δ*sun1 and* Δ*nek4* mutants produced some normal-sized oocysts that were able to sporulate, showing a partial rescue of the Δ*sun1* phenotype (**Fig. 3I**). In contrast, crosses between Δ*sun1* and Δ*hap2* did not rescue the Δ*sun1* phenotype. As controls, Δ*sun1,* Δ*hap2* and Δ*nek4* parasites alone were examined and no oocysts/sporozoites were detected (**Fig. 3I**). To further confirm the viability of controls, male (Δ*hap2)* and female (Δ*nek4)* mutants were crossed together and produced normal-sized oocysts that were able to sporulate (**Fig 3I**). These results indicate that a functional *sun1* gene is required from a male gamete for subsequent oocyst development. To assess the effect of the *sun1* deletion on DNA replication during male gametogenesis, we analysed the DNA content (N) of Δ*sun1* and WT-GFP male gametocytes by fluorometric analyses after DAPI staining. We observed that Δ*sun1* male gametocytes were octaploid (8N) at 8 min post activation, similar to WT-GFP parasites (**Fig 3I**), indicating that the absence of SUN1 had no effect on DNA replication. We also checked for the presence of DNA in gametes stained with Hoechst (a DNA dye) and found that most Δ*sun1* gametes were anucleate (**Fig 3J**).

### Transcriptomic and lipidomic analysis of **Δ***sun1* gametocytes shows no major changes in overall gene expression and lipid metabolism

To investigate the effect of *sun1* deletion on transcription, we performed RNA-seq analysis in triplicate on Δ*sun1* gametocytes and in duplicate on WT-GFP gametocytes, at 8 min post-activation. Total RNA was extracted to detect changes in gene expression post-activation. A detailed analysis of the mapped reads also confirmed the deletion of *sun1* in the Δ*sun1* strain (**Fig. S5D**). A relatively small number of differentially expressed genes was identified (**Fig. S5E**; **Supplementary Table S1**). Gene ontology (GO)-based enrichment analysis of these genes showed that several upregulated genes coded proteins involved in either lipid metabolic processes, microtubule cytoskeleton organization, or microtubule-based movement (**Fig. S5F**).

Given the nuclear membrane location of SUN1, we investigated the lipid profile of Δ*sun1* and WT-GFP gametocytes using mass spectrometry-based lipidomic analyses. Lipids are key elements of membrane biogenesis, with most phospholipids derived from fatty acid precursors (**Fig. S6A**).

We compared the lipid profile of Δ*sun1* and WT-GFP gametocytes, either non-activated (0 min) or activated for 8 min. Both the disruption of *sun1* and the activation of gametocytes caused minimal changes in the amounts of phosphatidic acid (PA), CE, monoacylglycerol (MAG), and TAG. Activated gametocytes had increased levels of PA, TAG, and CE and a decreased level of free fatty acids (FFA), compared to non-activated gametocytes (**Fig. S6B, C**). Following activation, there was also a differential level of lysophosphatidylcholine (LPC) and of specific fatty acid compositions (**Fig S6D**). Notably, the Δ*sun1* mutant had altered levels of arachidonic acid (C20:4), a scavenged fatty acid precursor of inflammatory molecules such as leukotrienes and prostaglandins (**Fig S6E**) (Botte et al., 2013; Dubois et al., 2018), and an increase in myristic acid (C14:0), a product of apicoplast fatty acid synthesis (FASII) (**Fig S6E**) (Amiar *et al*., 2020; Botte *et al*., 2013).

### Ultrastructural Microscopy Reveals Defects in Spindle Formation, Basal Body Segregation and Nuclear Attachment to Axonemes in **Δ***sun1* Gametocytes

To define the ultrastructural defects caused by *sun1* deletion, we used ultrastructure expansion microscopy (U-ExM) and transmission electron microscopy (TEM) to examine the morphology of male gametocytes at critical time points -1.5-, 8-, and 15-min post-activation, representing the timings of major transitions in spindle formation, BB segregation, and axoneme elongation (**Fig. 4A-D, Fig. S7, 8**).

**Fig 4.**
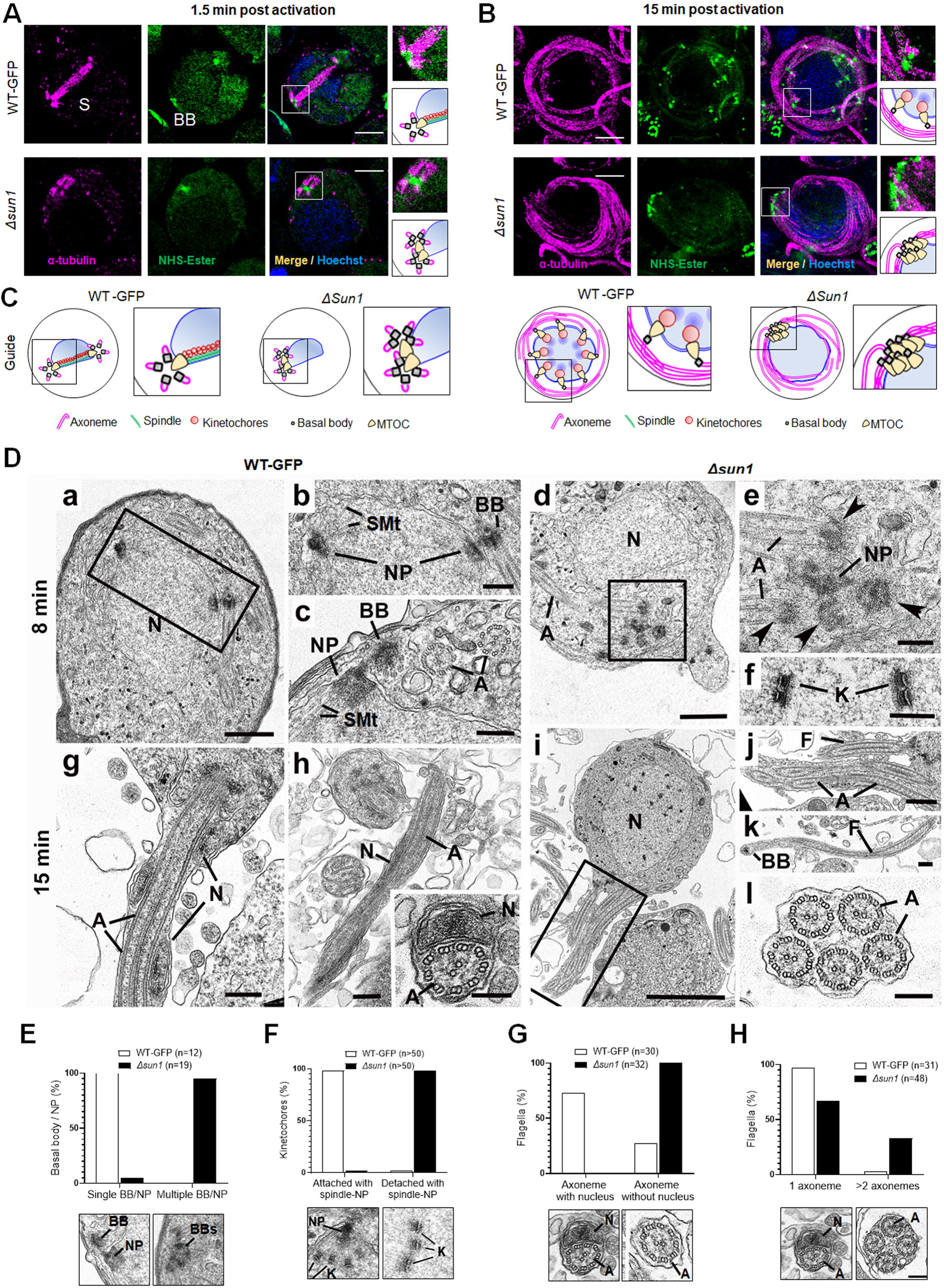
Ultrastructural analysis of Δ*sun1* gametocytes showing defect in spindle formation and BB segregation. **A.** Deletion of *sun*1 blocks first spindle formation as observed by ExM of gametocytes activated for 1.5 min. α -tubulin: magenta, amine groups/NHS-ester reactive: green. Basal Bodies: BB; spindle: S. Insets represent the zoomed area marked by the white boxes shown around BB/MTOC highlighted by NHS-ester and tubulin staining. Scale: 5LJµm. **B.** ExM images showing defect in BB/MTOC segregation in Δ*sun1* gametocytes activated for 15 min. α -tubulin: magenta, amine groups/NHS-ester reactive: green. Basal Bodies: BB; Insets represent the zoomed area shown around BB/MTOC highlighted by NHS-ester and tubulin staining. More than 30 images were analysed in more than three different experiments. Scale bars:LJ5LJµm. **C.** The schematic illustrates structures associated with mitosis and axoneme formation showing the first spindle is not formed, and BB are not separated in Δ*sun1* gametocytes. **D.** Electron micrographs of WT-GFP microgametocytes (male) at 8 min (**a-c**) and 15 min (**g, h**) plus Δ*sun1* gametocytes at 8 min (**d-f**) and 15 min (**i-l**). Bars represent 1 µm (**a, d, i**), and 100 nm (**b, c, e, f, g, h, insert, j, k, l**). **a.** Low power magnification of WT-GFP microgametocyte showing the central nucleus (N) with two nuclear poles (NP) with a basal body (BB) adjacent to one. The cytoplasm contains several axonemes (A). **b.** Enlargement of enclosed area (box) in **a** showing the basal body (BB) adjacent to one nuclear pole (NP). **c.** Detail showing the close relationship between the nuclear pole (NP) and the basal body (BB). Note the cross-sectioned axonemes (A) showing the 9+2 microtubular arrangement. **d.** Low power magnification of Δ*sun1* cell showing a cluster of electron dense basal structures (enclosed area) in the cytoplasm adjacent to the nucleus (N). A – axonemes. **e.** Detail from the cytoplasm (boxed area in **d**) shows a cluster of 4 basal bodies (arrowheads) and portions of axonemes (A) around a central electron dense structure of nuclear pole (NP) material. **f.** Detail from a nucleus showing kinetochores (K) with no attached microtubules. **g.** Periphery of a flagellating microgamete showing the flagellum and nucleus protruding from the microgametocyte. **h.** Detail of a longitudinal section of a microgamete showing the spiral relationship between the axoneme (A) and nucleus (N). **Insert.** Cross section of a microgamete showing 9+2 axoneme and adjacent nucleus (N). **i.** Section through a microgametocyte with a central nucleus (N) undergoing exflagellation. **j.** Enlargement of the enclosed area (box) in **i** showing one cytoplasmic protrusion containing a single axoneme forming a flagellum (F), while the other has multiple axonemes (A). **k.** Longitudinal section through a flagellum (F) with a basal body (B) at the anterior end but note the absence of a nucleus. **l.** Cross section showing a cytoplasmic process contain 5 axonemes (A) but no associated nucleus. **E-H.** Quantification of Δ*sun1* phenotypes compared to WT-GFP during male gametogenesis. N = Nucleus; BB = Basal Body; NP = Nuclear pole; A = Axonemes.

U-ExM revealed that in WT-GFP gametocytes, spindle microtubules were robustly assembled by 1.5 min post-activation, extending from nuclear MTOCs (spindle poles) to kinetochores. Simultaneously, BB - marked by NHS ester staining - segregated into tetrads, distributed across the cytoplasmic periphery (**Fig. 4A, S7A**). By 8 min, spindles were fully extended, ensuring accurate chromosome segregation, while BB had replicated to nucleate parallel axonemes aligned around the nucleus (**Fig. S7B**). By 15 min, these processes had culminated in exflagellation, producing mature microgametes containing nuclei tightly associated with BB-derived axonemes (**Fig. 4B, S7C**). In Δ*sun1* gametocytes, these processes were severely disrupted. At 1.5 min, α-tubulin staining showed incomplete or malformed spindles. NHS ester staining revealed BB clumped near one side of the nucleus, indicating a failure in segregation (**Fig. 4A, S7A**). By 8 min, spindles remained rudimentary, and BB segregation was still incomplete. Axoneme elongation proceeded, but BB and nuclear poles failed to align, leading to misconnected or unconnected axonemes (**Fig. S7B**). At 15 min, gametes had started emerging from cytoplasmic protrusions, but many lacked nuclei due to the absence of proper BB segregation and nuclear compartmentalization (**Fig. 4B**).

We performed TEM analysis of Δ*sun1* and WT-GFP gametocytes at 8- and 15 min post-activation. At 8 min post-activation, many of the wildtype male gametocytes had progressed to the third genome division with multiple nuclear spindle poles within a central nucleus (**Fig. 4Da; S8A**). In many cases, a BB was visible in the cytoplasm closely associated with and connected to the NP (**Fig. 4Db, c; S8A, B**). From the NP, nuclear spindle microtubules (SMt) radiated into the nucleoplasm and connected to kinetochores (**Fig. 4Db, c; S8A, B**). Within the cytoplasm several elongated axonemes, most but not all with the classical 9+2 microtubule arrangement, were visible (**Fig. 4Dc**).

At 8 min in Δ*sun1* gametocytes there were predominately mid stage forms with elongated axonemes running around the nucleus (**Fig. 4Dd; S8C, D, G**). It was also possible to find clumps (groups) of BB (**Fig. 4Dd, e; S8E, F**). Electron densities similar to NPs were observed adjacent to the BB (**Fig. 3De; S8H, I, J**). An extensive search (100+ sections) failed to identify examples of nuclear spindle formation with attached kinetochores. In contrast to the WT-GFP cells, a significant proportion of Δ*sun1* cells (20%) had groups of centrally located kinetochores with no attached microtubules (naked kinetochores: NK) (**Fig. 4Df; S8C, D, G**). NK were not observed in an extensive search of WT-GFP microgametocytes. (**Fig. S8A**). Nevertheless, the parallel orientation of the axonemes appeared to be maintained and many displayed the normal 9+2 arrangement (**Fig. 4Dc**).

By 15 min, WT-GFP gametocytes showed examples of exflagellation, with flagella displaying BB at the tip. Cross sections showed gametes with normal axonemes with a 9+2 arrangement and an enclosed nucleus surrounded by a plasma membrane (**Fig. 4Dg, h**). In Δ*sun1* samples, several late stages as well as some mid stages were observed. The late stages had more electron dense cytoplasm (**Fig. 4Di**). The majority had few if any axonemes, but all had a large nucleus with small clumps of electron dense material (**Fig. 4Di**). A few microgametocytes appeared to be undergoing exflagellation with flagellar-like structures protruding from the cell surface, often with multiple axonemes (**Fig. 4Dj, l**). However, they were abnormal, lacking a nucleus and often with variable numbers (1 to 6) of normal 9+2 axonemes (**Fig. 4Dj, k, l**). There appeared to be a disconnect between the axonemes and the nucleus during exflagellation. There was evidence of chromatin condensation to form microgamete nuclei but no connection between the nucleus and axoneme during exflagellation, resulting in microgametes lacking a nucleus. This Δ*sun1* mutant appeared to have a problem with separation of NPs and nuclear spindle formation, resulting in no genome segregation. The nuclear poles were difficult to find and did not appear to have been able to divide or move apart (**Fig. 4Dd, e; S8E, F, H, I, J**). It is possible that this lack of spindle pole division and movement was responsible for the BB remaining clumped together in the cytoplasm. It is also possible that clumping of the BB may explain the presence of multiple axonemes associated with the exflagellated structures. The quantification data for the Δ*sun1* phenotypes are shown in **Fig. 4E-H** and **Fig. S8K**.

Together, these findings highlight the pivotal role of SUN1 in coordinating the nuclear and cytoplasmic compartments during male gametogenesis. Its absence disrupts spindle formation, BB segregation, and the physical attachment of axonemes to the nucleus, resulting in anucleate microgametes incapable of fertilization.

### SUN1 Interactome Reveals Associations with Nuclear Envelope, ER and Chromatin Components

To identify protein interaction partners of PbSUN1, we performed GFP-based immunoprecipitation (IP) using a nanobody targeting SUN1-GFP in lysates of purified gametocytes activated for 6 to 8 min. This time point was chosen as it coincides with peak nuclear expansion and axoneme formation. To stabilize transient or weak interactions, protein cross-linking with 1% paraformaldehyde was used prior to IP. Co-immunoprecipitated proteins were identified using LC-MS/MS of tryptic peptides and were analysed using principal component analysis for duplicate WT-GFP and SUN1-GFP precipitations. (**Fig. 5A, Supplementary Table S2**).

**Fig. 5.**
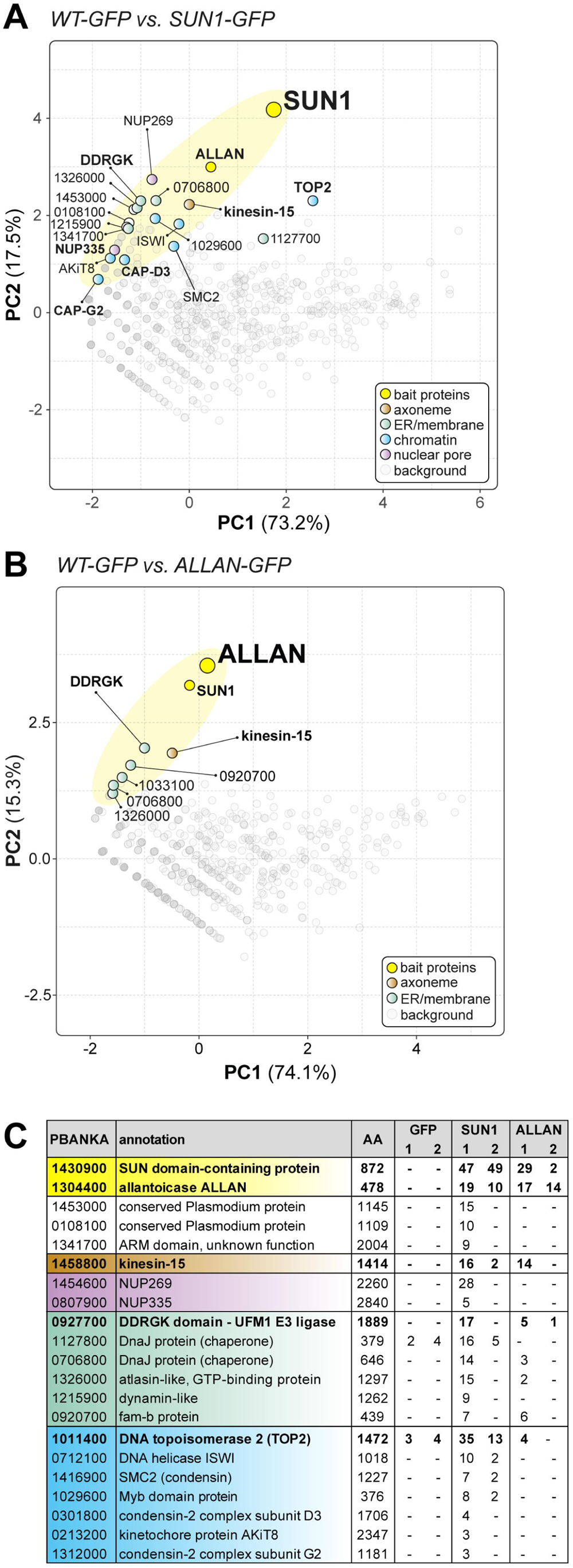
Reciprocal co-immunoprecipitation of PbSUN1-GFP and ALLAN-GFP during male gametogony. **A.** Projection of the first two components of a principal component analysis (PCA) of unique peptides derived from two SUN1-GFP (and WT-GFP) immunoprecipitations with GFP-trap (peptide values: **Supplementary Table S2**). A subset of proteins is highlighted on the map based on relevant functional categories. **B.** Similar to panel A, but now for the allantoicase-like protein ALLAN (PBANKA_1304400). **C.** Selected proteins, their size and corresponding gene ID and representation by the number of peptides in either WT-GFP, PbSUN1-GFP or ALLAN-GFP precipitates.

Co-variation with SUN1 was found for proteins of the nuclear envelope, chromatin and ER-related membranes. Notably, SUN1 co-purified with nuclear pore proteins (NUP269, NUP335), membrane proteins, including a likely ER component (DDRGK-domain containing UFM1 E3 ligase, PBANKA_0927700), chromatin-related factors (e.g. condensin I/II subunits, topoisomerase II and the kinetochore subunit AKit-8), and the cytoplasmic and male gametocyte-specific kinesin-15 (PBANKA_145880). This suggests a role for SUN1 in bridging chromatin (through condensin II) on the nuclear side and the cytoskeleton (through kinesin-15) on the cytoplasmic side of the nuclear envelope, potentially in association with nuclear pores (**Fig. 5A**) in a similar fashion to what has been proposed in the plant Arabidopsis (Ito et al., 2024; Sakamoto et al., 2022).

PbSUN1 also interacted with proteins harbouring a divergent carbohydrate-binding domain (like SUN1 itself), such as the allantoicase-like protein ALCC1 (Sayers et al. 2024), here referred to as ALLAN (PBANKA_1144200) and PBANKA_0209200, an ER-Golgi protein with a mannose-binding domain (**Fig. 5A, Supplementary Table S2**). ALLAN is an uncharacterized orthologue of allantoicase, an enzyme in purine metabolism. We could detect no KASH-like or lamin-like proteins in the co-immunoprecipitates. To further explore the interaction between ALLAN and SUN1, we performed the reciprocal ALLAN-GFP, from lysates of gametocytes 6-min post activation. Amongst the interactors we found SUN1 and its interactors, DDRGK-domain containing protein and kinesin-15 (**Fig. 5B, Supplementary Table S2**). The results from the immunoprecipitation experiments suggest that *Plasmodium* SUN1 functions in a non-canonical fashion via an interaction with ALLAN to tether chromatin to the nuclear envelope and possibly to the cytoplasmic cytoskeleton via kinesin-15 (**Fig. 5C**).

We hypothesized that SUN1 is at the centre of multiple molecular interactions during male gametogenesis, coordinating NE remodelling, spindle organization, and BB/axoneme attachment, with ALLAN as a key interactor near the nuclear MTOC.

### ALLAN is Located at Nuclear Poles (NP) and influences Basal Body/NP Segregation in Male Gametogenesis

To reveal ALLAN’s function in the *Plasmodium* life cycle, we generated a C-terminal GFP-tagged ALLAN parasite line, confirming correct integration by PCR (**Fig. S9A, B**) and detecting a protein of ∼85 kDa by Western blot (**Fig. S9C**). ALLAN-GFP parasites displayed normal growth and completed the life cycle, indicating that GFP tagging did not impair protein function.

During asexual blood stages, ALLAN-GFP exhibited a diffuse nucleoplasmic signal in trophozoites and schizonts with distinct focal points adjacent to dividing DNA, consistent with a role in mitotic regulation (**Fig. S9D**). However, by late schizogony, the ALLAN signal had diminished. In male and female gametocytes (**Fig. S9E**) and during zygote to ookinete transformation (**Fig. S9F**), ALLAN-GFP showed a spherical distribution around Hoechst-stained nuclear DNA and was enriched to form distinct focal points. During oocyst development and liver schizogony, ALLAN-GFP also exhibited distinct focal points adjacent to dividing DNA (**Fig. S9G, H**) suggesting a role in mitotic regulation, like in asexual blood stages (**Fig. S9D**).

In activated male gametocytes, ALLAN-GFP rapidly localized within a min post-activation to the NE, forming strong focal points that correlated with spindle poles or MTOCs (**Fig. 6A, Supplementary Video S2**). This localization persisted as ploidy increased from 1N to 8N in successive rounds of genome replication (**Fig. 6A**). Using U-ExM, ALLAN-GFP was resolved at the INM, enriched at spindle poles marked by NHS-ester staining, but absent from BB, where axonemes form (**Fig. 6B, C**). The location of ALLAN-GFP showed no overlap with that of kinesin-8B, a BB marker, or that of kinetochore marker NDC80 (**Fig. 6D, F**). Conversely, crosses with EB1-mCherry, a spindle marker, revealed an overlap at spindle poles (**Fig. 6E**) suggesting a role for ALLAN in nuclear spindle pole organization. Thus, the spatial relationship of SUN1 and ALLAN suggests coordinated roles in nuclear architecture and division of labour. SUN1 likely spans the NE, linking nuclear and cytoplasmic compartments, while ALLAN localizes more specifically to nuclear MTOCs.

**Fig. 6.**
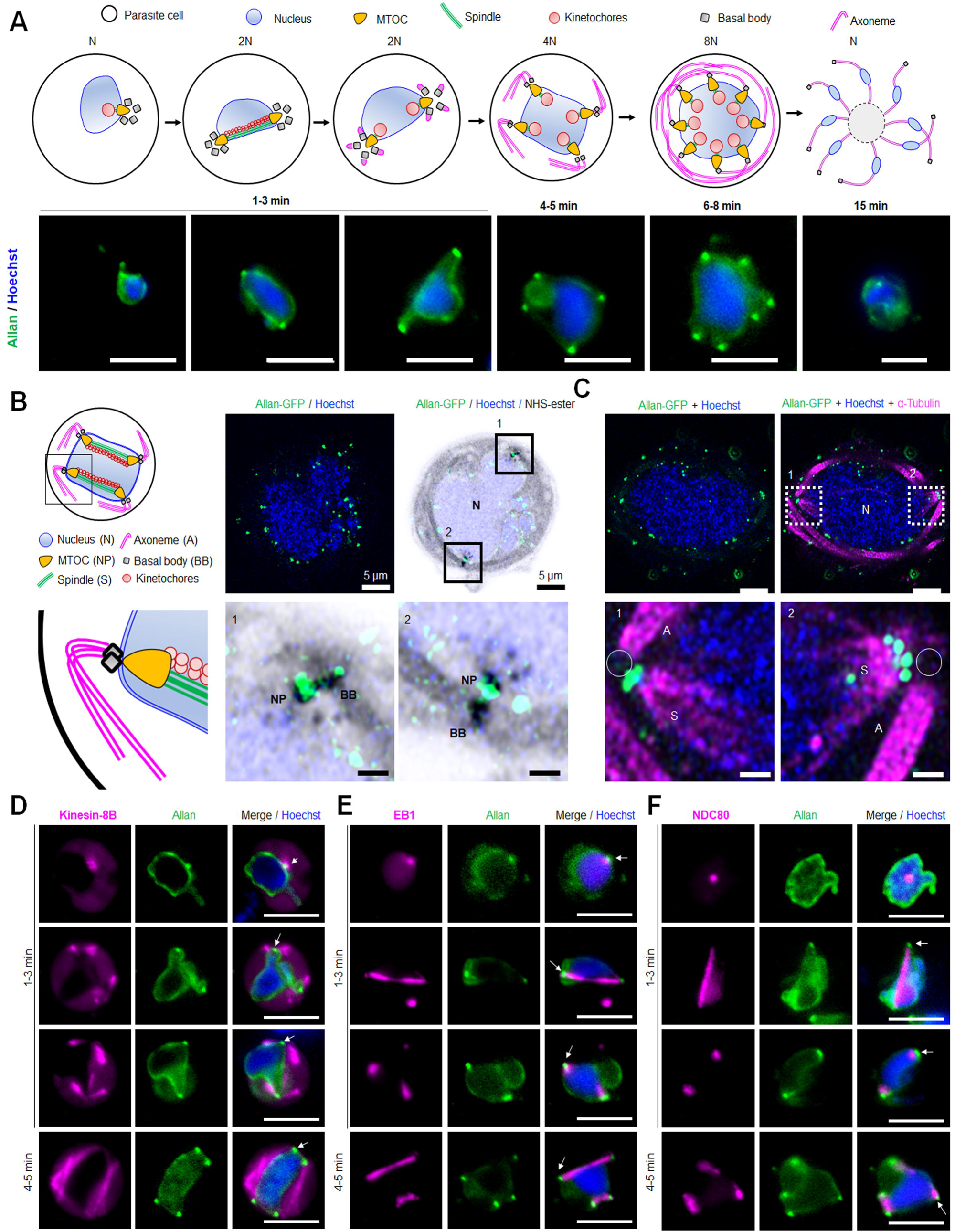
Location of ALLAN-GFP during male gametogenesis. **A.** The schematic on the upper panel illustrates the process of male gametogenesis. N, ploidy of nucleus. Live cell images showing the location of ALLAN-GFP (green) at different time points(1-15min) during male gametogenesis. Representative images of more than 50 cells with more than three biological replicates. Scale bar: 5 µm. **B.** ExM images showing location of ALLAN-GFP (green) detected by anti-GFP antibody compared to nuclear pole (NP)/MTOC and BB stained with NHS ester (grey) in gametocytes activated for 8 min. Scale bar: 5 µm. Representative images of more than 20 cells from two biological replicates. Insets represent the zoomed area shown around NP/MTOC and BB highlighted by NHS-ester. Scale bar: 1 µm. **C.** ExM images showing the location of ALLAN-GFP (green) compared to spindle and axonemes (magenta) detected by anti-GFP and antI-tubulin staining respectively in gametocytes activated for 8 min. Representative images of more than 20 cells from two biological replicates. Scale: 5 µm. Insets represent the zoomed area shown around spindle /axonemes highlighted by tubulin and GFP staining. Basal Bodies: BB; Spindle: S; Axonemes: A; Nucleus: N. Scale bar: 1 µm. **D, E, F.** Live cell imaging showing location of ALLAN-GFP (green) in relation to the BB and axoneme marker, kinesin-8B-mCherry (magenta) (D); spindle marker, EB1-mCherry (magenta) (E); and kinetochore marker, NDC80-mCherry (magenta) (F) during first mitotic division (1-3 min) of male gametogenesis. Arrows indicate the focal points of ALLAN-GFP. Representative images of more than 20 cells with three biological replicates. Scale bar: 5LJµm.

The proximity of both SUN1 and ALLAN at spindle poles is consistent with their collaboration to align spindle microtubules with nuclear and cytoplasmic MTOCs and ensure that chromosomal segregation aligns with axoneme formation. We performed functional studies of ALLAN deletion mutants to examine its contribution to spindle dynamics and its potential as a key player in rapid *Plasmodium* closed mitosis.

### Functional Role of ALLAN in Male Gametogenesis

We generated Δ*allan* mutants using double-crossover homologous recombination. Correct integration and successful gene deletion were confirmed via PCR (**Fig. S10A, S10B**) and qRT-PCR, with no residual *allan* transcription in Δ*allan* lines (**Fig. S10C**). There was no phenotype in asexual blood stages, but Δ*allan* mutants exhibited significant defects in mosquito stages.

Δ*allan* parasites showed no marked reduction in male gamete exflagellation compared to WT-GFP controls (**Fig. 7A**), and ookinete conversion rates were only slightly reduced (**Fig. 7B**). However, oocyst counts were significantly lower in Δ*allan*-parasite infected mosquitoes (**Fig 7C**), with diminished oocyst size (**Fig S10D**) and significant decrease in sporozoite number at 14-and 21-days post-infection (**Fig. 7D**). Though salivary gland sporozoites were significantly decreased (**Fig S10E**), Δ*allan* parasites were successfully transmitted to mice in bite-back experiments, showing that some viable sporozoites were produced (**Fig. 7E**).

**Fig. 7.**
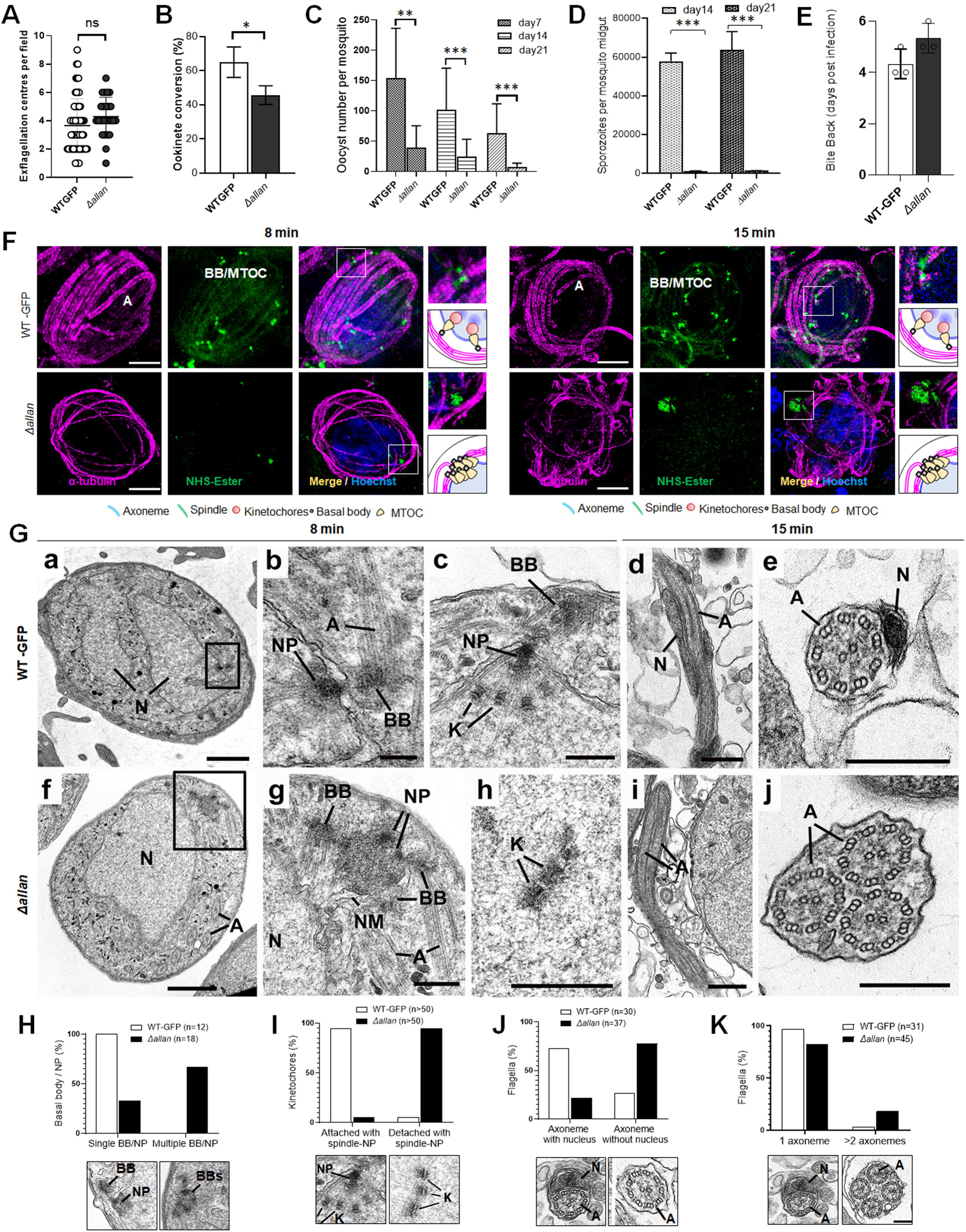
Deletion of ALLAN impairs male gametogenesis by blocking BB segregation. **A**. Exflagellation centres per field at 15 min post-activation in Δ*allan* compared to WT-GFP parasites. n≥3 independent experiments (>10 fields per experiment). Error bar, ± SEM. **B.** Percentage ookinete conversion from zygote. n≥3 independent experiments (> 100 cells). Error bar, ± SEM. **C.** Total number of GFP-positive oocysts per infected mosquito in Δ*allan* compared to WT-GFP parasites at 7-, 14-and 21-days post infection. Mean ± SEM. n≥3 independent experiments. **D.** Total number of sporozoites in oocysts of Δ*allan* compared to WT-GFP parasites at 14- and 21-days post infection. Mean ± SEM. n≥3 independent experiments. **E.** Bite back experiments reveal successful transmission of Δ*allan* and WT-GFP parasites from mosquito to mouse. Mean ± SEM. *n* = 3 independent experiments. **F.** ExM images of gametocytes activated for 8- and 15-min showing MTOC/BB stained with NHS ester (green) and axonemes stained with anti-tubulin antibody (magenta). Axonemes: A; Basal Bodies: BB; Microtubule organising centre: MTOC. Insets represent the zoomed area shown around BB/MTOC highlighted by NHS-ester and tubulin staining. More than 30 images were analysed in more than three different experiments. Scale bars:LJ5LJµm. **G.** Electron micrographs of WT-GFP microgametocytes at 8 mins (a-c) and 15 mins (d, e) and the Δ*allan* at 8 min (f-h) and 15 min (I, j). Bars represent 1 µm (a, f) and 200 nm in all other images. **a.** Low power image of a microgametocyte showing the nucleus (N) with two NP complexes (arrows) consisting of the basal body, NP and attached kinetochores. Axonemes (A) are present in the cytoplasm. **b.** Enlargement showing the nuclear pole (NP), associated basal body (BB) and axonemes (A). **c.** Detail of the nuclear pole (NP) showing kinetochores (K) attached the spindle microtubules. Note the basal body (BB) adjacent to the nuclear pole (NP). **d.** Longitudinal section of a microgamete showing the nucleus (N) closely associated with the axonemes (A). **e.** Cross section through a microgamete showing the nucleus (N) and axoneme (A) enclosed plasma membrane. **f.** Lower magnification of a microgametocyte showing a nucleus (N) with an adjacent clump of electron dense structures (enclosed area) and axonemes (A) in the cytoplasm. **g.** Enlargement of the enclosed area in f showing multiple basal bodies (BB) and unseparated nuclear poles (NP) enclosed by portions of nuclear membrane (NM). N – nucleus. **h.** Detail from a nucleus showing several kinetochores (K) with no associated spindle microtubules. **i.** Longitudinal section of an exflagellating cytoplasmic process consisting of two axonemes (A) but no nucleus. **j.** Cross section through an exflagellating cytoplasmic process showing the presence of multiple axonemes (A) but the absence of any nucleus. **H to K.** Quantification of Δ*allan* phenotype compared to WT-GFP during male gametogenesis. N = Nucleus; BB = Basal Body; NP = Nuclear pole; A = Axonemes.

U-ExM analysis revealed a striking defect in Δ*allan* male gametocyte morphology. At 8 min post-activation, NHS-ester staining indicated clustered BB, with incomplete segregation and misalignment relative to the nuclear MTOCs (**Fig. 7F**). Despite normal axoneme elongation, spindle organization was disrupted, as seen in nonsegregated BB/MTOCs (**Fig. 7F**).

To further explore the structural changes, we used electron microscopy analysis of gametocytes fixed at 8 min and 15 min post activation (**Fig. 7G**). At 8 min in the WT parasite there was a large lobated nucleus exhibiting multiple NPs. Radiating from the NPs were microtubules forming the nuclear spindle with associated kinetochores. BB was closely associated with the NP on the cytoplasmic side from which extended an axoneme (**Fig. 7Ga, b, c**). By 15 min, exflagellation had occurred and free microgametes were observed consisting of a 9+2 axoneme with closely associate nucleus enclosed by a unit membrane (plasmalemma) (**Fig. 7Gd, e**).

In the Δ*allan* parasites at 8 min, numerous mid stage microgametocytes were observed. However, in contrast to the WT parasites, the BB appeared to be clumped together although axoneme formation was similar to that in WT. (**Fig. 7Gf, g**). NP-like structures lacking spindle microtubules were often associated with clumps of electron dense material (**Fig. 7Gg; S11D, E**). Examination of sections of mid stage gametocytes (50+), revealed several cells (∼20%) with free kinetochores and no associated spindle microtubules (**Fig. 6Gh; S11A, B, C**). It was difficult to find individual NPs or obvious nuclear spindles although very rare examples were observed (>2%).

By 15 min post-activation in Δ*allan* gametocytes, several cells had exflagellated with the formation of numerous free gametes – the majority with no associated nucleus (**Fig. 7Gi; S11G, H**). Some had multiple axonemes (**Fig. 7Gi, j; S11H**) but there were many fewer examples than seen in the Δ*sun1* mutant. However, unlike the Δ*sun1* line, a few male gametes with axonemes and associated nuclei were observed. The Δ*allan* phenotype quantitative data are shown in **Fig. 7H-K** and **Fig. S11I**.

These Δ*allan* defects suggest that ALLAN is important for the separation of the NPs with formation of spindle microtubules and kinetochore attachment. The low incidence of spindle assembly and impaired NP organization in Δ*allan* gametocytes is reminiscent of the Δ*sun1* mutant phenotype, reinforcing the relationship between these two proteins. The presence of a few normal nuclear spindle poles with attached kinetochores and formation of a few microgametes with axoneme and nucleus may explain the transmission observed for this mutant.

### Evolution of a Novel SUN1-ALLAN Axis in Haemosporida

The non-canonical SUN1-ALLAN complex in *P. berghei* is a unique apicomplexan adaptation, where NE and MTOC dynamics diverge from those of other eukaryotes. To trace the evolution of this interaction, we examined SUN1, ALLAN, and potential interactors (e.g., KASH proteins and lamins) across alveolates (apicomplexans, dinoflagellates and ciliates) and model organisms such as yeast, humans, and *Arabidopsis* (**Fig. 8, Supplementary Table S3**).

**Fig 8.**
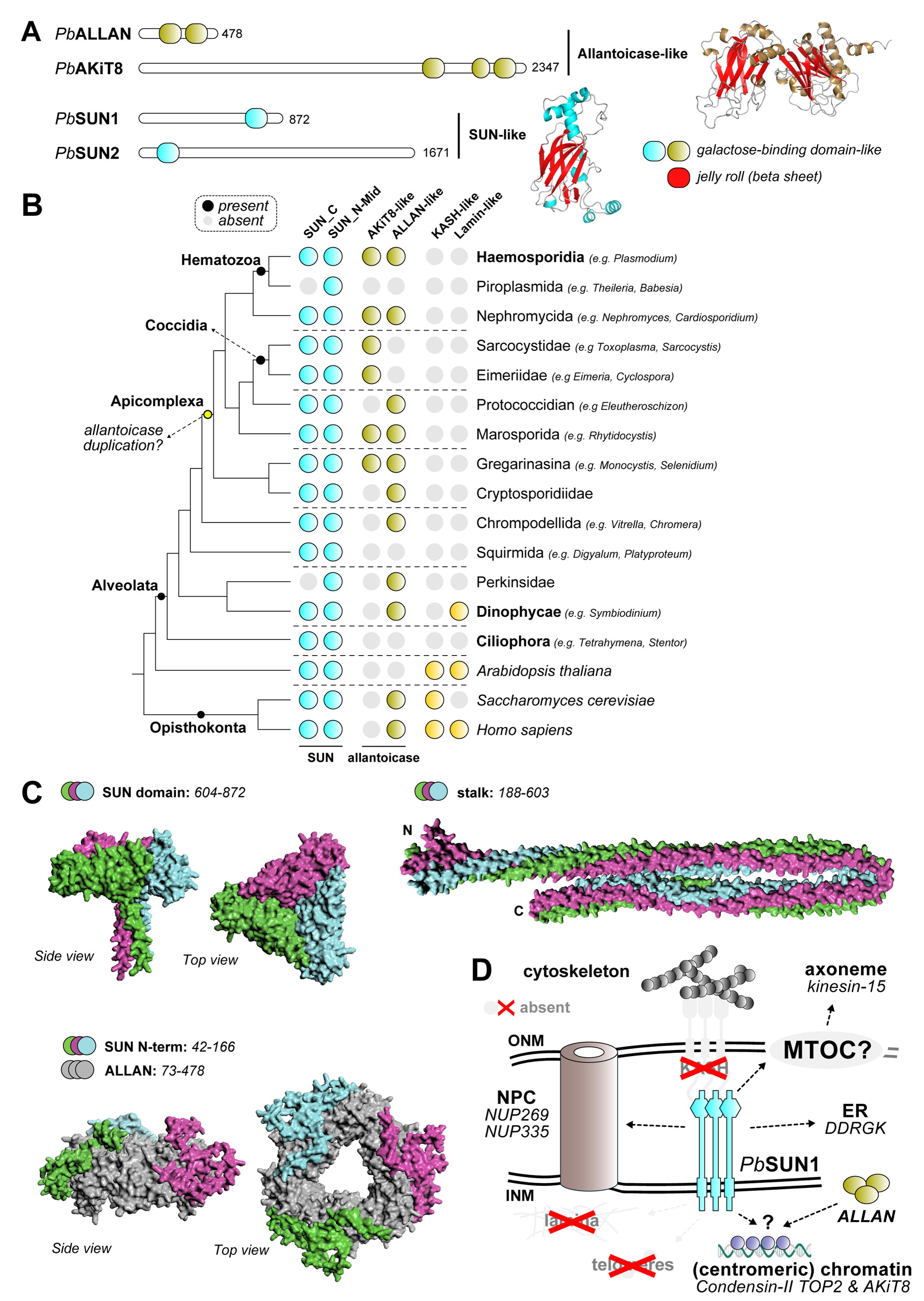
Evolution and Structure of the SUN1-ALLAN interaction. **A.** Domain analysis shows two proteins with allantoicase domain and two proteins with SUN-domain in *P. berghei.* The SUN domain and two domains comprising allantoicases are part of the same galactose-binding domain family, with a strikingly similar fold. **B.** Phylogenetic profiles showing the presence of SUN-, ALLAN- and KASH-domain and lamin proteins in Apicomplexa and a selection of other eukaryotes, including two model species *Homo sapiens* and *Arabidopsis thaliana.* **C.** AlphaFold3-modelled interaction between ALLAN and SUN1 based on separate domains (no full structure could be modelled). The SUN1 C-terminus forms a trimeric complex (pTM:0.37) similar to a trimeric ALLAN complex (grey) with the N-terminus of SUN1 interacting with ALLAN (pTM:0.55). This N-terminal domain is unique to Haemosporida. **D.** Overview in similar style as Fig. 1 of main interactors for putative localization at the nuclear envelope for ALLAN and SUN1 during male gametogenesis. Structures in grey have not been found to be associated with SUN1.

SUN domain proteins generally fall into two families: C-terminal SUN domain and N-terminal/mid SUN domain proteins (Graumann et al., 2014). Most eukaryotes have representatives of both families. *P. berghei* has two SUN domain proteins (Kandelis-Shalev et al, 2024), one of each family with PbSUN1 having a C-terminal SUN domain. Allantoicase-like proteins often consist of two nearly identical domains, belonging to the galactose-binding-like family that are remarkably similar to the SUN domain, (**Fig. 8A**). Among apicomplexans, two allantoicase subtypes are present due to duplication in the ancestor of all Apicomplexa: AKiT8 (Brusini et al., 2022) a kinetochore-associated protein, and ALLAN, the allantoicase-like subtype. Notably, *Plasmodium* is one of the few apicomplexans retaining both subtypes (i.e. only AKiT is retained in Toxoplasma). Extreme sequence divergence complicates phylogenetic classification of allantoicase-like proteins among apicomplexans, but length-based classification distinguished the longer AKiT8 (∼2300 aa) from ALLAN-like proteins (∼800 aa; **Fig. 8B**). Despite using previously developed Hidden Markov Model and structural searches (Benz et al., 2024), we found no evidence of KASH-domain proteins or lamins in amongst apicomplexans, similar to what was reported in prior studies (**Fig. 8B**) (Koreny and Field, 2016).

AlphaFold3 (AF3) modelling suggested that SUN1 and ALLAN interact specifically in Haemosporida through the N-terminal domain of SUN1, a region that is absent from other apicomplexan and/or other eukaryotic SUN1 family proteins (**Fig. 8C**). We could not detect similar interactions between SUN1-like and ALLAN-like orthologs amongst other apicomplexans, ruling out a rapidly evolving interaction. Based on these structural predictions and the super-resolution imaging of SUN1-and ALLAN-GFP lines (**Fig. 2D, E**; **Fig. 6B, C**), we propose that ALLAN resides on the nucleoplasmic side of the NE, while SUN1’s C-terminal domain likely extends toward the ONM to interact with proteins involved in axoneme and cytoskeletal dynamics in the nucleoplasm (**Fig. 8D**). This specificity may underscore the restricted nature of a SUN1-ALLAN complex to Haemosporida, and thus specific adaption of the allantoicase-like protein ALLAN for nuclear envelope dynamics.

## Discussion

This study identifies the SUN1-ALLAN complex as a novel and essential mediator of NE remodelling and bipartite MTOC coordination during *P. berghei* male gametogenesis. *Plasmodium* lacks KASH-domain proteins and lamins, and a canonical LINC complex, relying on a highly divergent mechanism to tether nuclear and cytoplasmic compartments during rapid closed mitosis (Rout et al., 2017).

Our results reveal that SUN1 and ALLAN form a unique complex essential for spindle assembly, BB segregation, and axoneme organization, maintaining an intact NE throughout. Disruption of either protein causes spindle and MTOC miscoordination, leading to defective male gametes incapable of fertilization. Additionally, SUN1 deletion alters lipid homeostasis and NE dynamics, underscoring its multifaceted role in nuclear organization and metabolism. These findings establish the SUN1-ALLAN axis as a crucial evolutionary innovation in *Plasmodium*.

In *P. berghei* male gametogenesis, mitosis is closed and extraordinarily rapid with features that diverge significantly from those of classical models. Within just 8 min, there are three rounds of DNA replication (1N to 8N), accompanied by the assembly of axonemes on cytoplasmic BB (Guttery *et al*., 2022). KASH-domain proteins and lamins are absent, yet *Plasmodium* achieves precise nuclear and cytoskeletal coordination. Our findings reveal that this is accomplished through a non-canonical SUN1-ALLAN complex. SUN1 located at the NE, forms loops and folds to accommodate rapid nuclear expansion, and ALLAN facilitate spindle pole coordination.

Recent studies on SUN1-domain containing proteins have been reported in some Apicomplexa parasites including *Toxoplasma* and *Plasmodium* (Kandelis-Shalev et al., 2024; Sayers et al., 2024; Wagner et al., 2023). SUN-like protein-1 of *Toxoplasma gondii* shows a mitotic spindle pole (MTOC) location and is important for nuclear division (Wagner *et al*., 2023). *Plasmodium spp.* encode two SUN proteins (SUN1 and SUN2), that show a nuclear membrane location and play roles in nuclear division and DNA repair during the blood stage of *P. falciparum* (Kandelis-Shalev *et al*., 2024). Recent work by Sayers et al. (2024) demonstrated that PbSUN1-HA was associated with the NE and suggested that it may be required to capture the intranuclear MTOC onto the NE (Sayers *et al*., 2024). They also showed that it forms a complex with ALCC1 (the protein we have designated ALLAN). They showed that SUN1 is essential for fertile male gamete formation and that in SUN1-KO lines basal bodies are aggregated. Our study extends their findings by providing live-cell imaging of SUN1 and ALLAN and their coordination with key mitotic markers such as NDC80, EB1, and Kinesin-8B. We generated and characterised both SUN1 and ALLAN knockout parasites, performing ultrastructural analyses using TEM and extensive functional studies. Consistent with the results of Sayers et al., we observed basal body clumping in SUN1-KO, and showed that kinetochore attachment to the spindle is compromised in both mutant parasites. Our work emphasizes the role of the nuclear envelope in the rapid cell division during male gametogenesis. Collectively, the findings from both studies highlight the importance of the SUN1–ALLAN complex in Plasmodium cell division (Sayers *et al*., 2024).

*Plasmodium* SUN1 shares some similarities with SUN proteins in other organisms like *S. cerevisiae and Schizosaccharomyces pombe* (Fan et al., 2022; Hagan and Yanagida, 1995), but its interaction with ALLAN is a highly specialized adaptation in *Plasmodium*. The location of SUN1 at the nuclear MTOC, distinct from axonemal markers like kinesin-8B, underscores its role in nuclear compartmentalization. Similarly, the absence of functional spindle formation and kinetochore attachment in Δ*sun1* mutants is reminiscent of phenotypes seen in mutants of spindle-associated proteins in *Plasmodium*, such as EB1, ARK2 and kinesin-8X (Zeeshan *et al*., 2023; Zeeshan et al., 2019b). However, SUN1’s role extends specifically to NE remodelling and the organization of the inner acentriolar MTOC.

Our study demonstrates that SUN1 is primarily located in the space between the INM and ONM, with the NE but facing the nucleoplasm via it’s N-terminus, forming dynamic loops and folds to accommodate the rapid expansion of the nucleus during three rounds of genome replication. These loops serve as structural hubs for spindle assembly and kinetochore attachment at the nuclear MTOC, separating nuclear and cytoplasmic compartments. The absence of SUN1 disrupts spindle formation, causing defects in chromosome segregation and kinetochore attachment, while BB segregation remains incomplete, resulting in clumped basal bodies and anuclear flagellated gametes. This phenotype highlights the role of SUN1 in maintaining the nuclear compartment of the bipartite MTOC.

ALLAN, a novel allantoicase-like protein, has a location complementary to that of SUN1, forming focal points at the inner side of the NE. Like SUN1, ALLAN is also not essential during asexual erythrocytic stage proliferation. Functional studies following ALLAN gene deletion revealed similar defects in MTOC organization, with impaired spindle formation, kinetochore attachment, and nuclear-cytoplasmic coordination during flagellated gamete assembly. These findings indicate that SUN1 and ALLAN work together, replacing the role of lamins and KASH proteins in coordinating nuclear and cytoskeletal dynamics.

Interestingly, our interactome analysis identified kinesin-15 as a putative interactor of SUN1 and ALLAN, suggesting a possible link between this motor protein and nuclear remodelling. Previous studies have shown that kinesin-15 is essential for male gamete formation and is located partially at the plasma membrane and nuclear periphery, albeit mostly cytoplasmic (Zeeshan et al., 2022b). On the nuclear side, we found subunits of the condensin-II complex, similar to what was found in Arabidopsis (Ito *et al*., 2024; Sakamoto *et al*., 2022). These interactions with SUN1 and ALLAN raise the possibility of a broader network of proteins facilitating nuclear and cytoskeletal coordination during male gametogenesis.

Transcriptomic analysis of the Δ*sun1* mutant showed upregulation of genes involved in lipid metabolism and microtubule organization. Lipidomic profiling further confirmed substantial alterations in the lipid composition of SUN1-deficient gametocytes. Notably, PA and CE, both critical for membrane curvature and expansion, were reduced in the Δ*sun1* mutant, whereas MAG and DAG levels were elevated. These disruptions likely reflect impaired NE expansion and structural integrity during gametogenesis. Additionally, the altered levels of specific fatty acids, such as arachidonic acid and myristic acid, suggest perturbations in both scavenged and apicoplast-derived lipid pathways..

The SUN1-ALLAN complex exemplifies a lineage-specific adaptation in *P. berghei*, highlighting the evolutionary plasticity of NE remodelling mechanisms across eukaryotes. SUN-domain proteins, a hallmark of LINC complexes, are widely conserved and diversified across eukaryotic lineages, yet *Plasmodium* and related apicomplexans lack key canonical LINC components.

Our analyses suggest that the SUN1-ALLAN complex is specific to Haemosporida. While SUN-domain proteins are conserved across most eukaryotes, the allantoicase-like protein ALLAN is a novel addition to the nuclear architecture of apicomplexans. Phylogenetic profiling indicates that ALLAN likely arose from a duplication of an allantoicase in an early apicomplexan ancestor, likely the ancestor to all coccidians and hematozoa. This event gave rise to two subtypes: AKiT8, associated with kinetochores (Brusini *et al*., 2022), and ALLAN, specialized in NE dynamics.

The absence of KASH proteins and lamins in apicomplexans raises fascinating questions about how NE and cytoskeletal coordination evolved in this group. Comparative studies in ichthyosporeans, close relatives of fungi and animals, have identified intermediate forms of mitosis involving partial NE disassembly (Shah et al., 2024). In contrast, *Plasmodium* employs a highly streamlined closed mitosis, likely driven by the requirement for rapid nuclear and cytoskeletal coordination during male gametogenesis. The discovery of the SUN1-ALLAN axis suggests that apicomplexans have developed alternative strategies to adapt the NE for their cellular and mitotic requirements. These findings raise intriguing questions about the evolutionary pressures that shaped this unique nuclear-cytoskeletal interface in apicomplexans. The presence of similar or other non-canonical complexes in other organisms remains an open question, with potential implications for understanding the diversity of mitotic processes.

This study opens several avenues for future research into the biology of the malaria parasite and its unique adaptations for mitosis. While the SUN1-ALLAN complex has been shown to play a central role in bipartite MTOC organisation during male gametogenesis, many questions remain about its broader role and potential applications. Another key question concerns the evolutionary origins of the SUN1-ALLAN axis; comparative studies across apicomplexans, dinoflagellates, ciliates, and other non-model eukaryotes may help trace the evolution of this complex. The presence of similar non-canonical complexes in other rapidly dividing eukaryotic systems may shed light on the diversity of mitotic adaptations beyond *Plasmodium*. A functional further exploration of associated proteins, such as kinesin-15, may help elucidate how the SUN1-ALLAN complex integrates with other cellular machinery and structures. Investigating whether kinesin-15, a motor protein, interacts directly with SUN1 or ALLAN and has a catalytic role in nuclear-cytoskeletal remodelling may be critical to further our understand of how the system achieves such rapid mitotic coordination.

## Methods

### Ethics statement and mice details

The animal work passed an ethical review process and was approved by the United Kingdom Home Office. Work was carried out under UK Home Office Project Licenses (PDD2D5182 and PP3589958) in accordance with the UK ‘Animals (Scientific Procedures) Act 1986’. Six-to eight-week-old female CD1 outbred mice from Charles River laboratories were used for all experiments. The mice were maintained under a 12 h light and 12 h dark (7 am till 7LJpm) cycle, at a temperature between 20 and 24 °C, and a humidity between 40 and 60%.

### Generation of transgenic parasites and genotype analyses

To generate lines for GFP-tagged SUN1 and ALLAN, a region of each gene downstream of the ATG start codon was amplified, ligated to the p277 vector, and transfected as previously described (Guttery et al., 2014). The p277 vector includes a human DHFR cassette, providing resistance to pyrimethamine. Schematic representations of the endogenous gene loci, the vector constructs, and the recombined gene loci can be found in **Supplementary Figs. 1A** and 6A. For parasites expressing C-terminal GFP-tagged proteins, diagnostic PCR was performed with primer 1 (Int primer) and primer 2 (ol492) to confirm integration of the GFP - targeting constructs (**Supplementary Figs. 1B and 6B**). The primers used to generate the mutant parasite lines can be found in Supplementary Data 3.

Gene-deletion targeting vectors for SUN1, and ALLAN were created using the pBS-DHFR plasmid. This plasmid contains polylinker sites flanking a *Toxoplasma gondii* dhfr/ts expression cassette, which provides resistance to pyrimethamine, as described previously (Saini et al., 2017). PCR primers N1511 and N1512 were used to amplify a 1,1094 bp fragment of the 5′ sequence upstream of *sun1* from genomic DNA, which was then inserted into the ApaI and HindIII restriction sites upstream of the dhfr/ts cassette in the pBS-DHFR plasmid. A 776 bp fragment from the 3′ flanking region of *sun1* was generated with primers N1513 and N1514, and inserted downstream of the dhfr/ts cassette using the EcoRI and XbaI restriction sites. The same approach was used for *allan,* amplifying upstream (1044 bp) and downstream (1034 bp) sequences and insertion into the pBS-DHFR plasmid. The linear targeting sequence was released from the plasmid using *Apa*I/*Xba*I digestion. A schematic representation of the endogenous *sun1* and *allan* loci, the construct and the recombined *sun1* and *allan* loci are presented in Supplementary Fig. 1A and 6A, respectively. The primers used to generate these mutant parasite lines can be found in **Supplementary Table S4**. A diagnostic PCR used primer 1 (IntN151_5) and primer 2 (ol248) to confirm integration of the targeting construct, and primer 3 (Int303) and primer 4 (N1514) were used to confirm deletion of the *sun1* gene (**Supplementary Fig. S1B** and **Supplementary Table S4**). Similarly, a diagnostic PCR using primer 1 (IntN153_5) and primer 2 (ol248) was used to confirm integration of the targeting construct, and primer 3 (Int307) and primer 4 (N1534) were used to confirm deletion of the *allan* gene (**Supplementary Fig. S6B and Supplementary Table S4**). *P. berghei* ANKA line 2.34 (for GFP-tagging), and ANKA line 507cl1 expressing GFP (for the gene-deletion) were transfected by electroporation (Janse *et al*., 2006).

### Purification of schizonts and gametocytes

Blood cells obtained from infected mice (day 4 post infection) were cultured for 11 h and 24 h at 37°C (with rotation at 100 rpm) and schizonts were purified the following day on a 60% v/v NycoDenz (in phosphate-buffered saline (PBS)) gradient, (NycoDenz stock solution: 27.6% w/v NycoDenz in 5 mM Tris-HCl (pH 7.20), 3 mM KCl, 0.3 mM EDTA).

Gametocytes were purified using a modified version of the protocol of Beetsma et al. (1998). In brief, parasites were injected into mice pre-treated with phenylhydrazine and enriched by sulfadiazine treatment two days post-infection. Blood was collected on the fourth day post-infection, and gametocyte-infected cells were purified using a 48% NycoDenz gradient (prepared with NycoDenz stock solution as above). Gametocytes were harvested from the gradient interface and washed thoroughly.

### Live-cell imaging

To investigate SUN1-GFP, and ALLAN-GFP expression during erythrocytic stages, parasites cultured in schizont medium were imaged at various stages of schizogony. Purified gametocytes were assessed for GFP expression and location at different time points (0 and 1 to 15 min) post-activation in ookinete medium (RPMI 1640 medium containing 100 μM xanthurenic acid, 1% w/v sodium bicarbonate, and 20 % v/v heat inactivated foetal bovine serum [FBS]). Zygote and ookinete stages were labelled using the cy3-conjugated 13.1 antibody (red), which targets the P28 surface protein, and examined over a 24-hour period. Oocysts and sporozoites were imaged in infected mosquito guts. All images were captured with a 63× oil immersion objective on a Zeiss AxioImager M2 microscope equipped with an AxioCam ICc1 digital camera.

### Western blot analysis

Purified gametocytes were placed in lysis buffer (10 mM Tris-HCl [pH 7.5], 150 mM NaCl, 0.5 mM EDTA, and 1% NP-40). The lysed samples were placed for 10 min at 95°C after adding Laemmli buffer, and then centrifuged at 13,000 g for 5 min. Samples were electrophoresed on a 4% to 12% SDS-polyacrylamide gel, and resolved proteins were transferred to nitrocellulose membrane (Amersham Biosciences). Immunoblotting was performed using the Western Breeze Chemiluminescence Anti-Rabbit kit (Invitrogen) and an anti-GFP polyclonal antibody (Invitrogen) at a dilution of 1:1,250, according to the manufacturer’s instructions.

### Generation of dual tagged parasite lines

The GFP (green)-tagged SUN1 or ALLAN parasites were mixed in equal numbers with mCherry (red)-tagged lines of kinetochore marker (NDC80), basal body/axoneme marker (kinesin-8B), and spindle markers (EB1 and ARK2) and injected into mice. Mosquitoes fed on these mice 4 to 5 days post-infection, when gametocytaemia was high, were monitored for oocyst development and sporozoite formation at days 14 and 21 after feeding. Infected mosquitoes were then allowed to feed on naïve mice, and after 4 to 5 days, these mice were examined for blood-stage parasitaemia by microscopy of Giemsa-stained blood smears. Some parasites expressed both GFP- and mCherry-tagged proteins in the resultant gametocytes; these cells were purified, and fluorescence microscopy images were collected as described above.

### Parasite phenotype analyses

Blood samples containing approximately 50,000 SUN1-knockout, or ALLAN-knockout parasites were injected intraperitoneally (i.p.) into mice. Asexual stage parasite development and gametocyte production were monitored by microscopy on Giemsa-stained thin smears. Four to five days post-infection, gametocytes were harvested and exflagellation and ookinete conversion were examined as described above, using a Zeiss AxioImager M2 microscope. To analyze mosquito infection and parasite transmission, 30 to 50 *Anopheles stephensi* SD 500 mosquitoes were allowed to feed for 20 min on anesthetized, infected mice that had at least 15% asexual parasitaemia and a comparable gametocyte level. To assess midgut infection, approximately 15 guts were dissected from mosquitoes on days 7 and 14 post-feeding, and oocysts were counted by microscopy using a 63× oil immersion objective. On day 21 post-feeding, another 20 mosquitoes were dissected, and their guts and salivary glands were crushed separately in a loosely fitting homogenizer to release sporozoites, which were then quantified using a haemocytometer or used for imaging.

Mosquito bite-back experiments with naïve mice were conducted 21 days post-feeding, and blood smears were examined after 3 to 4 days.

### Immunoprecipitation and Mass Spectrometry

Purified male gametocyte pellets from SUN1-GFP, and ALLAN-GFP parasites at 6-min post-activation. were cross-linked by 10 min incubation with 1% formaldehyde, followed by 5 min incubation in 0.125 M glycine solution and three washes with phosphate-buffered saline [PBS; pH 7.5]. WT-GFP gametocytes were used as controls. Cell lysates were prepared and immunoprecipitation was conducted using a GFP-Trap_A Kit (Chromotek) according to the manufacturer’s instructions. Briefly, lysates were incubated for 2 hours with GFP-Trap_A beads at 4°C with continuous rotation, then unbound proteins were washed away, and bound proteins were digested with trypsin. The tryptic peptides were analyzed by liquid chromatography-tandem mass spectrometry. Mascot (http://www.matrixscience.com/) and MaxQuant (https://www.maxquant.org/) search engines were used for mass spectrometry data analysis. Experiments were performed in duplicate. Peptides and proteins with a minimum threshold of 95% were used for further proteomic analysis. The PlasmoDB database was used for protein annotation, and manual curation was used to classify proteins into categories relevant for SUN1/ALLAN interactions: chromatin, nuclear pore, ER membrane and axonemal proteins. To capture co-variation of bound proteins between different experiments (comparing GFP-only with SUN1-GFP and ALLAN-GFP) we performed a principal component analysis (PCA) using unique peptide values per protein present in two replicates per experiment. Values for undetected proteins were set to 0. Values were ln (xLJ+LJ1) transformed and PCA was performed using the ClustVis webserver (settings Nipals PCA, no scaling) (Metsalu and Vilo, 2015).

### AlphaFold3 modelling

3D protein structures for SUN1 and ALLAN were modelled using the AlphaFold3 webserver (https://alphafoldserver.com/) with standard settings (seed set to 100) (Abramson et al., 2024). To assess the stoichiometry of the complex we modelled from monomers up to decamers for both ALLAN and SUN1, using the N-terminus, the middle domain and the C-terminus of SUN1. ALLAN consistently formed higher order structures containing trimeric complexes. Although SUN1 could form higher order structures beyond trimers, we opted to model it as a trimer given the preference of ALLAN to form trimers.

### Fixed Immunofluorescence Assay

SUN1-KO and ALLAN-KO gametocytes were purified, activated in ookinete medium, fixed at various time points with 4% paraformaldehyde (PFA, Sigma) diluted in microtubule (MT)-stabilizing buffer (MTSB) for 10 to 15 min, and added to poly-L-lysine coated slides. Immunocytochemistry used mouse anti-α tubulin mAb (Sigma-T9026; used at 1:1000) as primary antibody, and secondary antibody was Alexa 568 conjugated anti-mouse IgG (Invitrogen-A11004) (used at 1:1000). Slides were mounted in Vectashield with DAPI (Vector Labs) for fluorescence microscopy with a Zeiss AxioImager M2 microscope fitted with an AxioCam ICc1 digital camera.

### Liver stage cultures

Human liver hepatocellular carcinoma (HepG2) cells (European Cell Culture Collection) were grown in DMEM medium, supplemented with 10% heat inactivated FBS, 0.2% NaHCO_3_, 1% sodium pyruvate, 1% penicillin/streptomycin and 1% L-glutamine in a humidified incubator at 37LJ°C with 5% CO_2_. For infection, 1LJ×LJ10^5^ HepG2 cells were seeded in a 48-well culture plate. The day after seeding, sporozoites were purified following mechanical disruption from salivary glands removed from female *A. stephensi* mosquitoes infected with PbALLAN-GFP parasites, and added in infection medium to the culture maintained for 72 h.

### Ultrastructure Expansion Microscopy (UExM)

Purified gametocytes were activated for different time periods, and then activation was stopped by adding 4% paraformaldehyde. Fixed cells were then attached to a 12LJmm diameter poly-D-lysine (A3890401, Gibco) coated coverslip for 10 min. Coverslips were incubated overnight in 1.4% formaldehyde (FA)/2% acrylamide (AA) at 4°C. Gelation was performed in ammonium persulfate (APS)/TEMED (10% each)/monomer solution (23% sodium acrylate; 10% AA; 0.1% BIS-AA in PBS) for 1 h at 37°C. Gels were denatured for 15 min at 37°C and 45 min at 95°C and then incubated in distilled water overnight for complete expansion. The following day, gels were washed twice in PBS for 15 min to remove excess water, and then incubated with primary antibodies at 37°C for 3 h. After washing 3 times for 10 min in PBS/ 0.1% Tween, incubation with secondary antibodies was performed for 3 h at 37°C followed by 3 washes of 10 min each in PBS/ 0.1% Tween (all antibody incubation steps were performed at 37°C with 120 to 160 rpm shaking). Directly after antibody staining, gels were incubated in 1 ml of 594 NHS-ester (Merck: 08741) diluted to 10 μg/ml in PBS for 90 min at room temperature on a shaker. The gels were then washed 3 times for 15 min with PBS/0.1% Tween and expanded overnight in ultrapure water. One cm × 1 cm pieces were cut from the expanded gel and attached to 24 mm diameter Poly-D-Lysine (A3890401, Gibco) coated coverslips to prevent the gel from sliding and avoid drifting while imaging. The primary antibody was either against α-tubulin (1:1000 dilution, Sigma-T9026), or an anti-GFP antibody (1:250, Thermo Fisher). Secondary antibodies were anti-rabbit Alexa 488, or anti-mouse Alexa 568 (Invitrogen), used at dilutions of 1:1000. Atto 594 NHS-ester (Merck 08741) was used for bulk proteome labelling. Images were acquired on Zeiss Elyra PS.1-LSM780 and CD7-LSM900, and Airyscan confocal microscopes, where 0.4 Airy unit (AU) on confocal and 0.2 AU were used along with slow scan modes. Image analysis was performed using Zeiss Zen 2012 Black edition and Fiji-Image J.

### Structured Illumination Microscopy

A small volume (3 µl) of gametocyte suspension was placed on a microscope slide and covered with a long (50 × 34 mm) coverslip to obtain a very thin monolayer and immobilize the cells. Cells were scanned with an inverted microscope using a Zeiss Plan-Apochromat 63x/1.4 oil immersion or Zeiss C-Apochromat 63x/1.2 W Korr M27 water immersion objective on a Zeiss Elyra PS.1 microscope, utilizing the structured illumination microscopy (SIM) technique. The correction collar of the objective was set to 0.17 for optimal contrast. The following settings were used in SIM mode: lasers, 405 nm: 20%, 488 nm: 16%; exposure times 200 ms (Hoechst), 100 ms (GFP), three grid rotations, five phases. The bandpass filters BP 420-480 + LP 750, and BP 495-550 + LP 750 were used for the blue and green channels, respectively. Where multiple focal planes (Z-stacks) were recorded processing and channel alignment was done as described previously (Zeeshan et al., 2024).

### Electron microscopy

Gametocytes activated for 8 min and 15 min were fixed in 4% glutaraldehyde in 0.1 M phosphate buffer and processed for electron microscopy (Ferguson et al., 2005). Briefly, samples were post fixed in osmium tetroxide, treated en bloc with uranyl acetate, dehydrated, and embedded in Spurr’s epoxy resin. Thin sections were stained with uranyl acetate and lead citrate prior to examination in a JEOL JEM-1400 electron microscope (JEOL, United Kingdom). The experiments were done at least 3 times to capture every stage of the parasite, and 50 to 55 cells were examined for phenotypic analysis. For more details, please see the figure legends.

### RNA Isolation and Quantitative Real-Time PCR (qRT-PCR) Analyses

RNA was extracted from purified gametocytes using an RNA purification kit (Stratagene). Complementary DNA (cDNA) was synthesized using an RNA-to-cDNA kit (Applied Biosystems). Gene expression was quantified from 80 ng of total RNA using the SYBR Green Fast Master Mix kit (Applied Biosystems). All primers were designed using Primer3 (Primer-BLAST, NCBI). The analysis was conducted on an Applied Biosystems 7500 Fast machine with the following cycling conditions: 95°C for 20 sec, followed by 40 cycles of 95°C for 3 sec and 60°C for 30 sec. Three technical replicates and three biological replicates were performed for each gene tested. The genes hsp70 (PBANKA_081890) and arginyl-tRNA synthetase (PBANKA_143420) were used as endogenous control reference genes. The primers used for qPCR are listed in **Supplementary Table S4**.

### RNA-seq Analysis

Libraries were prepared from lyophilized total RNA, starting with the isolation of mRNA using the NEBNext Poly(A) mRNA Magnetic Isolation Module (NEB), followed by the NEBNext Ultra Directional RNA Library Prep Kit (NEB) as per the manufacturer’s instructions. Libraries were amplified through 12 PCR cycles (12 cycles of [15 s at 98°C, 30 s at 55°C, 30 s at 62°C]) using the KAPA HiFi HotStart Ready Mix (KAPA Biosystems). Sequencing was performed on a NovaSeq 6000 DNA sequencer (Illumina), generating paired-end 100-bp reads.

FastQC [https://www.bioinformatics.babraham.ac.uk/projects/fastqc/ was used to analyze raw read quality and the adapter sequences were removed using Trimmomatic (v0.39) [http://www.usadellab.org/cms/?page=trimmomatic]. The resulting reads were mapped against the P. *berghei* genome (PlasmoDB, v68) using HISAT2 (v2-2.2.1) with the --very-sensitive parameter. Uniquely mapped, properly paired reads with mapping quality of 40 or higher were retained using SAMtools (v1.19) [http://samtools.sourceforge.net/]. Raw read counts were determined for each gene in the *P. berghei* genome using BedTools [https://bedtools.readthedocs.io/en/latest/ #] to intersect the aligned reads with the genome annotation. Differential expression analysis was performed using DESeq2 to call up-and down-regulated genes (FDR < 0.05 and log2 FC > 1.0). Volcano plots were made using the R package Enhanced Volcano.

### Statistical Analysis

All statistical analyses were conducted using GraphPad Prism 9 (GraphPad Software). Student’s t-test and/or a two-way ANOVA test were employed to assess differences between control and experimental groups. Statistical significance is indicated as *P < 0.05, **P < 0.01, ***P < 0.001, or ns for not significant. ’n’ denotes the sample size in each group or the number of biological replicates. For qRT-PCR data, a multiple comparisons t-test, with post hoc Holm–Sidak test, was utilized to evaluate significant differences between wild-type and mutant parasites.

## Data availability

The RNA-seq data generated in this study have been deposited in the NCBI Sequence Read Archive. Mass spectrometry proteomics data have been deposited to the ProteomeXchange Consortium via the PRIDE partner repository. Source data are provided with this paper.

## Acknowledgements

R.T. is supported by ERC advance grant funded by UKRI Frontier Science (EP/X024776/1), MRC UK (MR/K011782/1) and BBSRC (BB/L013827/1, BB/X014681/1). M.Z, I.B., D.B. A.M. were supported as research fellows by (EP/X024776/1). R.Y. is supported by BBSRC (BB/X014681/1). S.L.P is supported by Wellcome DBT India Alliance/Team Science (IA/TSG/21/1/600261). AAH is supported by the Francis Crick Institute (FC001097), which receives core funding from the Cancer Research UK (FC001097), the UK Medical Research Council (FC001097), and the Wellcome Trust (FC001097). KGLR is supported by the NIH/NIAID (R01 AI136511) and the University of California, Riverside (NIFA-Hatch-225935. E.T. was supported by a personal fellowship from the Nederlandse Organisatie voor Wetenschappelijk Onderzoek (NOW), the Netherlands (grant no. VI. Veni.202.223). YYB and CYB were supported by Agence Nationale de la Recherche, France (Project ApicoLipiAdapt grant ANR-21-CE44-0010 ; Project Apicolipidtraffic grant ANR-23-CE15-0009-01 ; Project OIL grant ANR-24-CE15-2171-02), The Fondation pour la Recherche Médicale (FRM EQU202103012700), Laboratoire d’Excellence Parafrap, France (grant ANR-11-LABX-0024), LIA-IRP CNRS Program (Apicolipid project), the Université Grenoble Alpes (IDEX ISP Apicolipid) and Région Auvergne Rhone-Alpes for the lipidomics analyses platform (Grant IRICE Project GEMELI), Collaborative Research Program Grant CEFIPRA (Project 6003-1) by the CEFIPRA (MESRI-DBT). Confocal and SIM microscopy was conducted in the School of Life Sciences Imaging (SLIM). For Open Access, the authors have applied a CC BY public copyright licence to any Author Accepted Manuscript version arising from this submission. We thank Cleidiane Zampronio at Warwick University for mass spectrometry methods and Bio Support Unit, University of Nottingham for maintenance of mice used in this study.

## Contributions

R.T. conceived and coordinated the project; M.Z., I.B. and R.T. performed and analysed the live-cell imaging, knockout, and proteomics data; M.Z., R.T., I.B. and D.B. performed mice and mosquito-related work; R.Y. and D.J.P.F performed and analysed electron microscopy imaging. M.Z., S.L.P. and R.Y. performed expansion microscopy; Z.C., B.M. and S.B. performed RNA-seq analysis; Y.Y.B. and C.Y.B. did lipid analysis; A.M. performed liver stage microscopy; M.H., D.J.P.F. and S.V. performed SBF-SEM and 3D nuclear modelling. R.M. performed structured illumination microscopy; I.B. and D.B. performed qRT PCRs; A.B. performed mass spectrometry analysis; E.C.T. performed the interactomics, structural and phylogenetic analyses; S. B. performed, structural analyses; R.T., and A.A.H. supervised the project; M.Z., D.B. and R.T. managed the project. M.Z. E.C.T. and R.T. prepared the first manuscript draft. A.A.H. helped significantly in editing and writing and all authors contributed to manuscript editing and revisions.

