## Supplementary figures and images for "A novel SUN1-ALLAN complex coordinates segregation of the bipartite MTOC across the nuclear envelope during rapid closed mitosis in Plasmodium"

### Fig S1

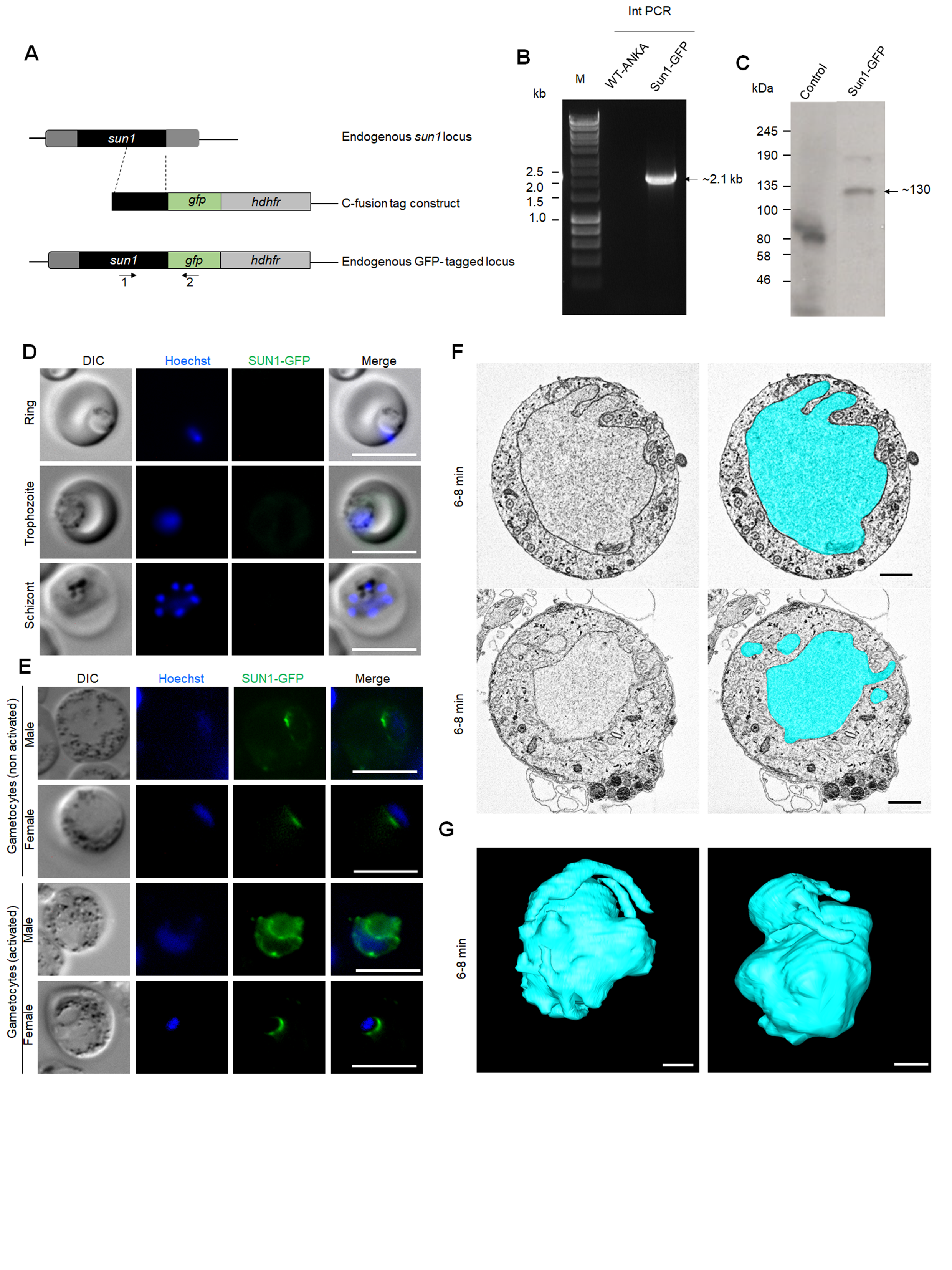

### Fig S2

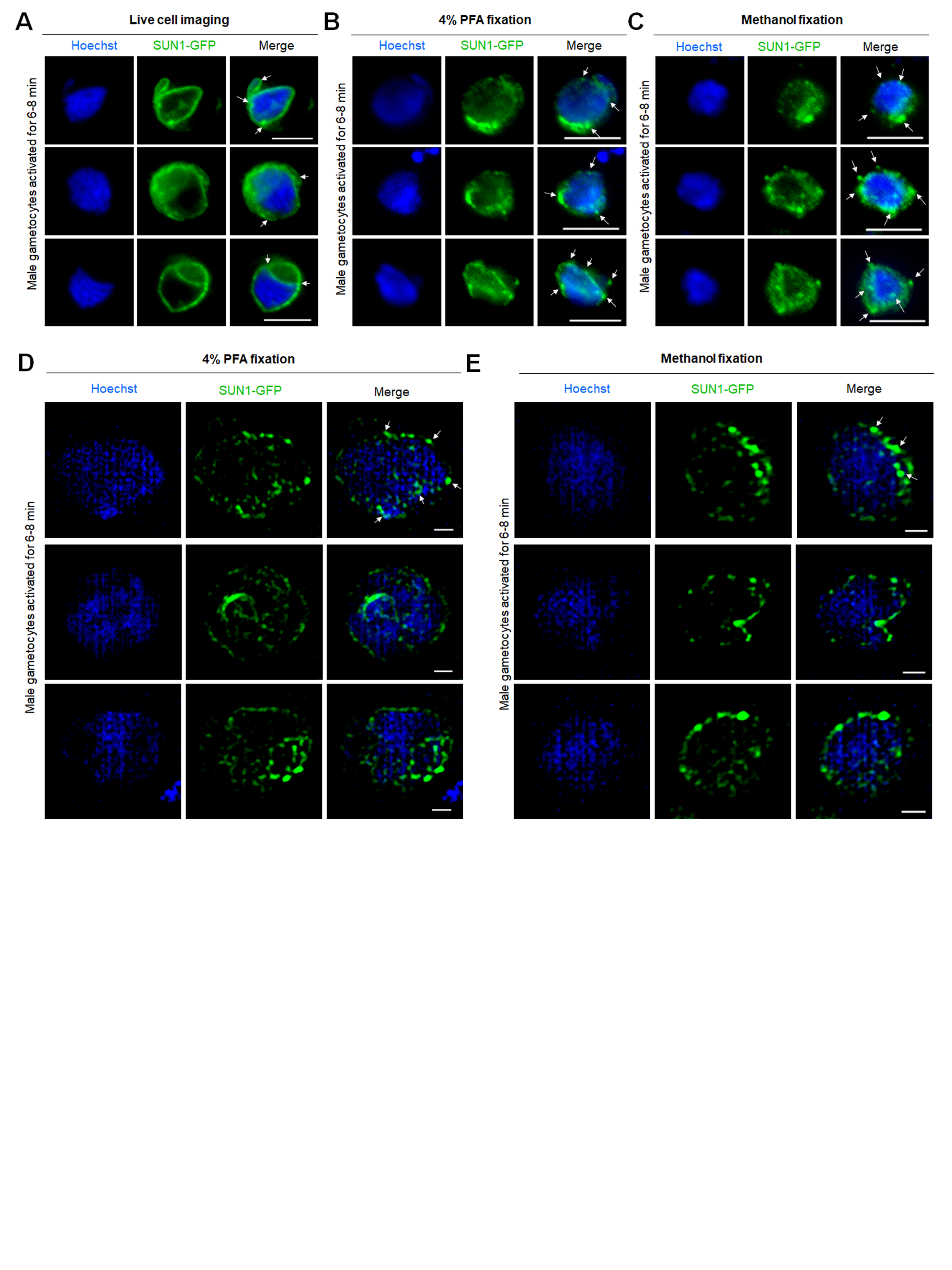

### Fig S3

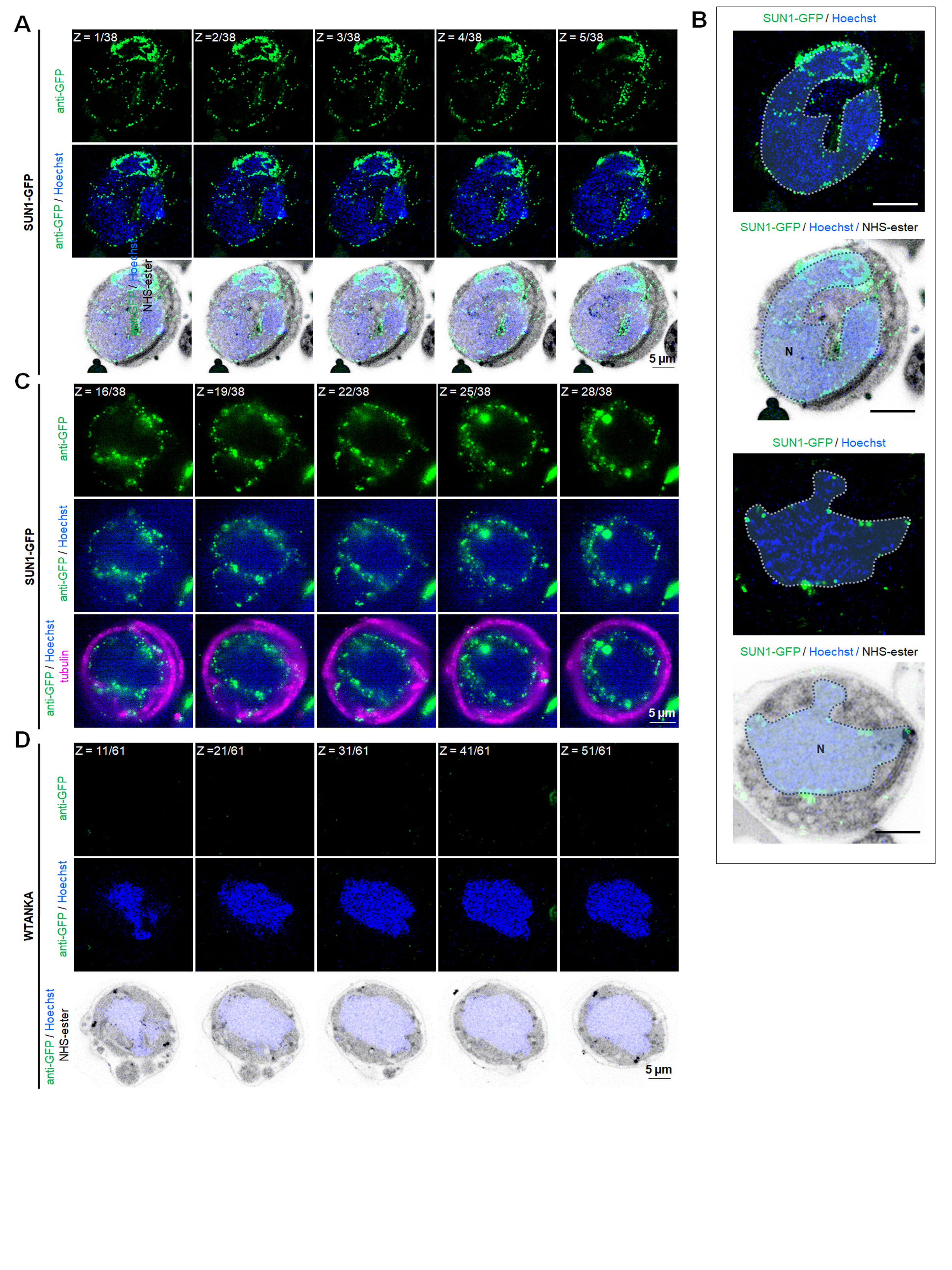

### Fig S4

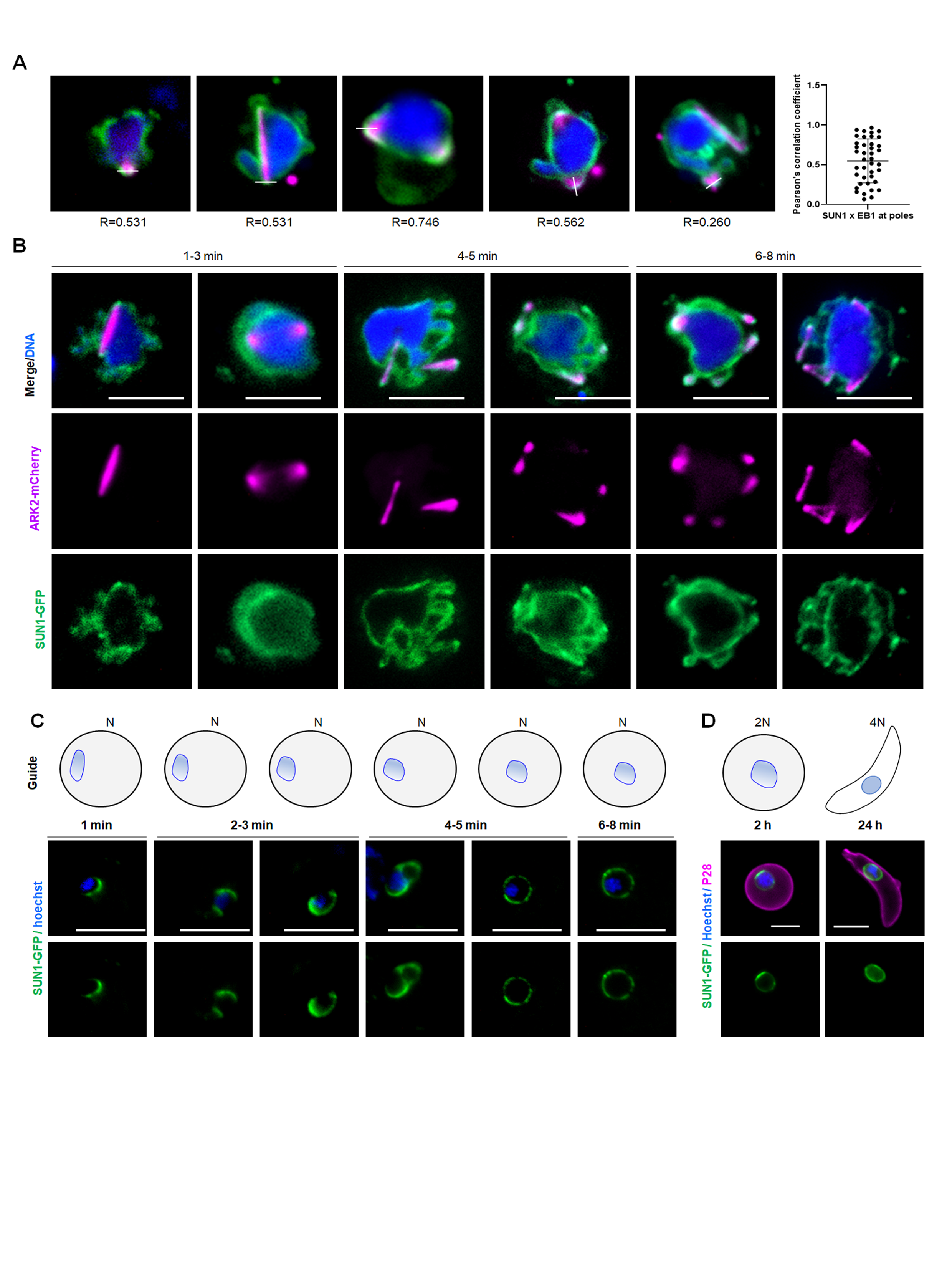

### Fig S5

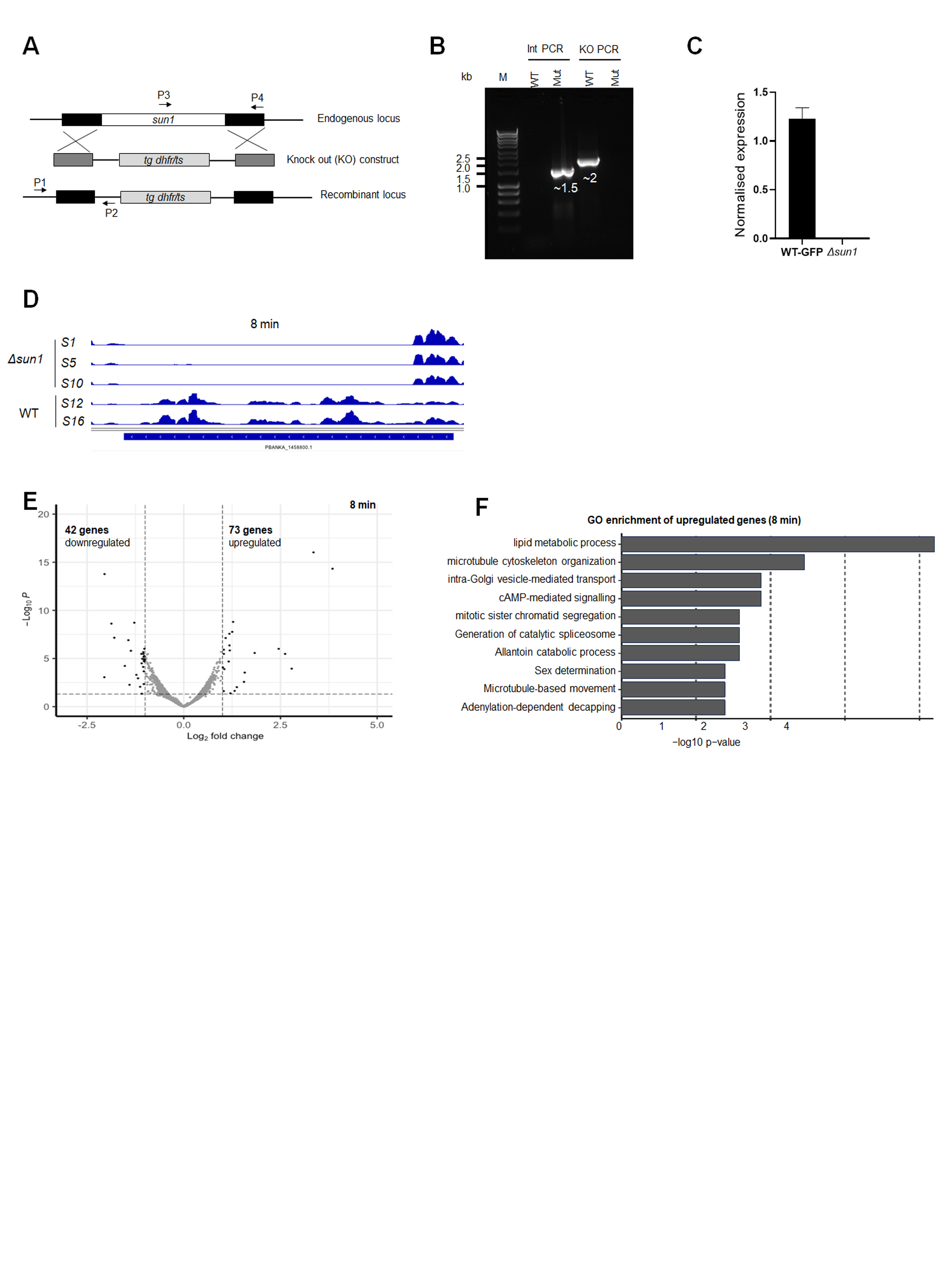

### Fig S6

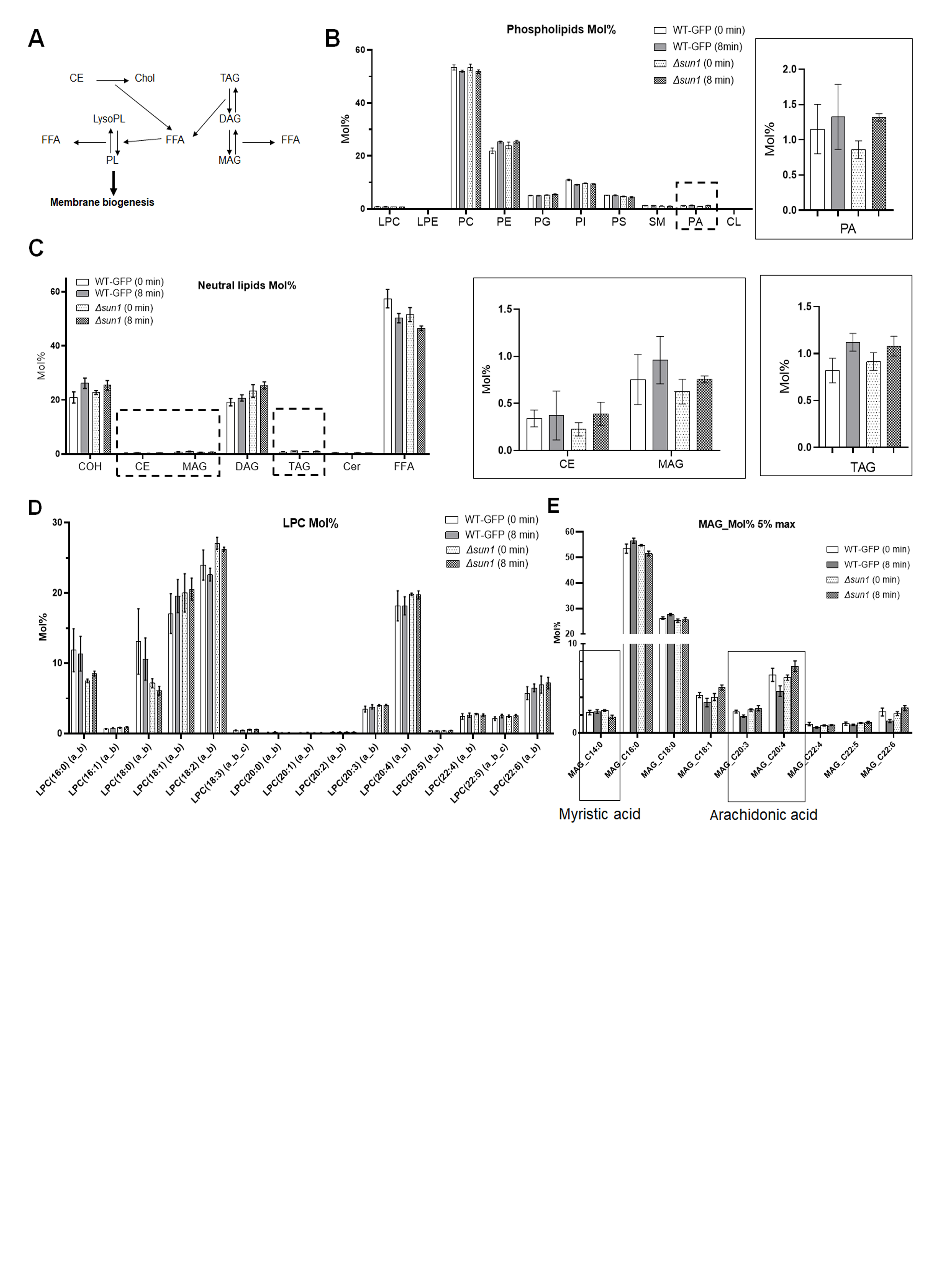

### Fig S7

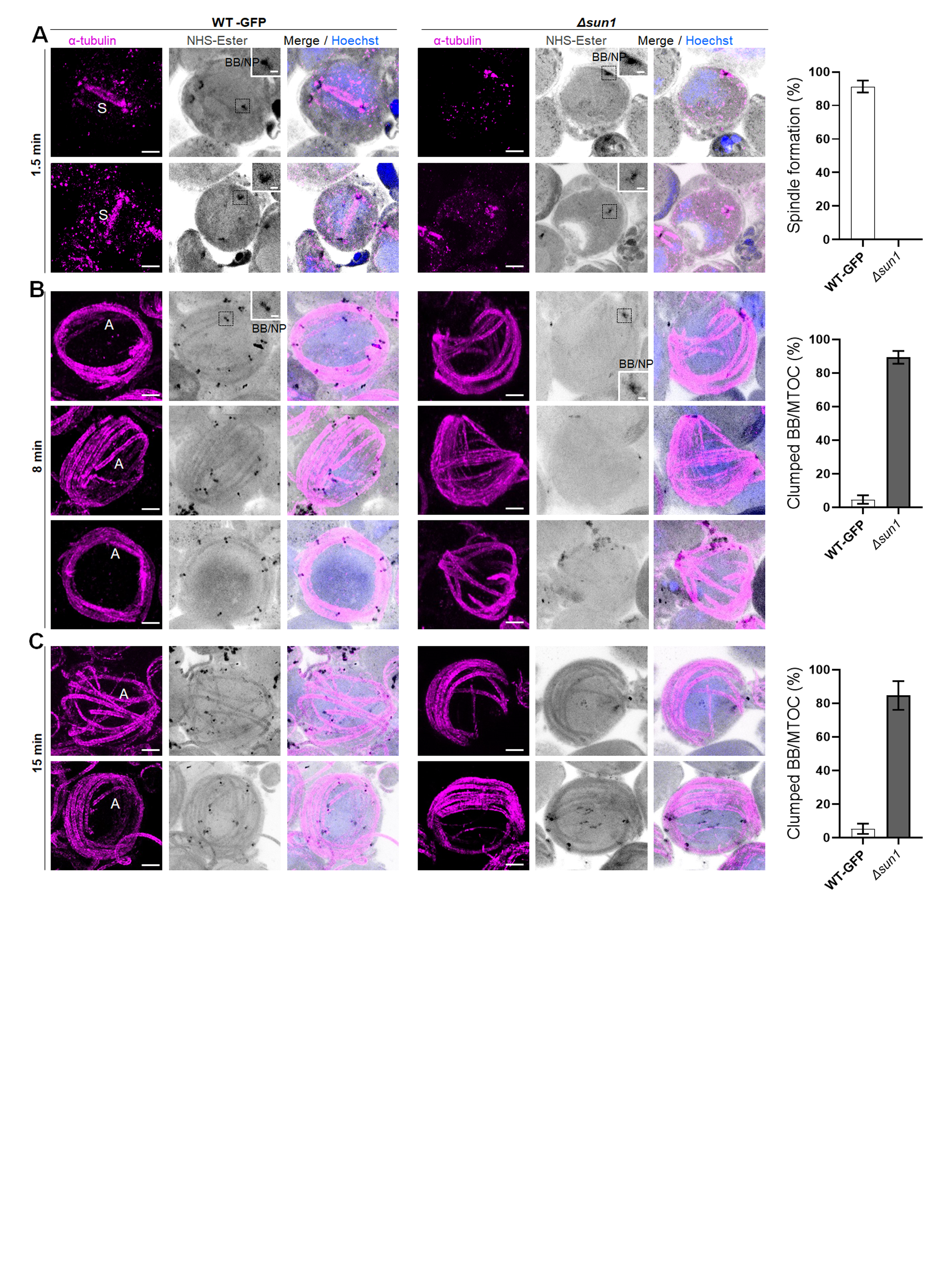

### Fig S8

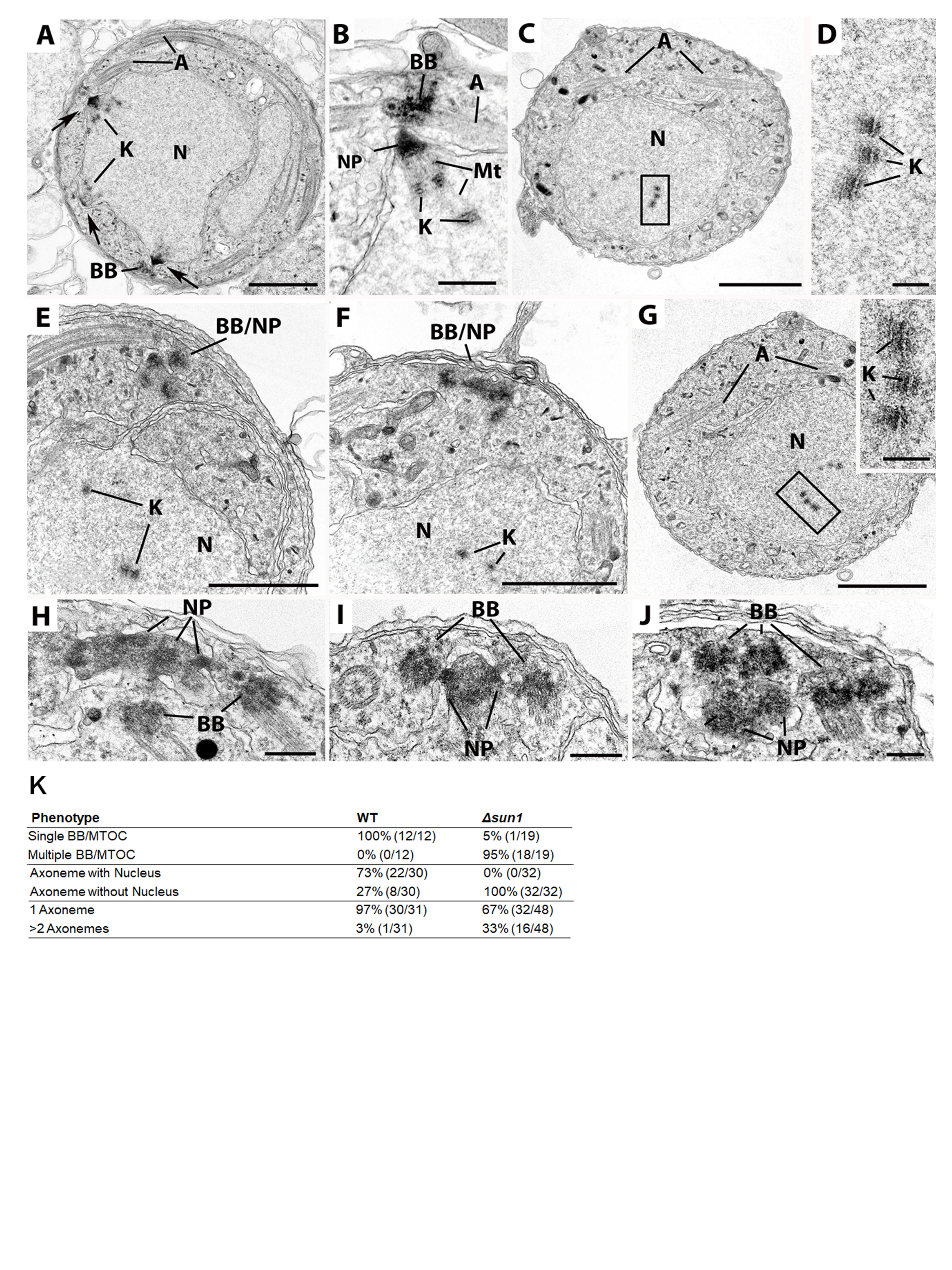

### Fig S9

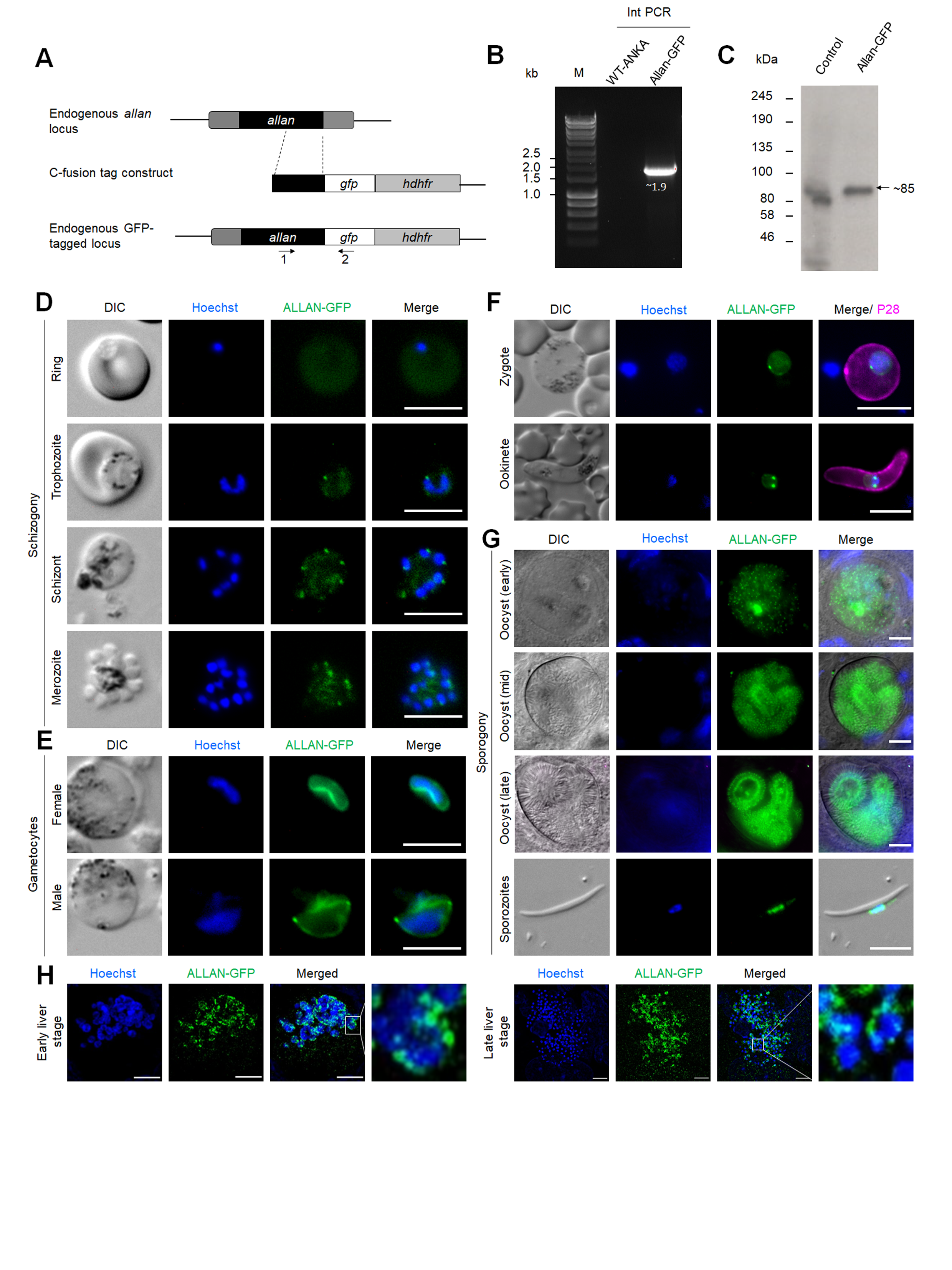

### Fig S10

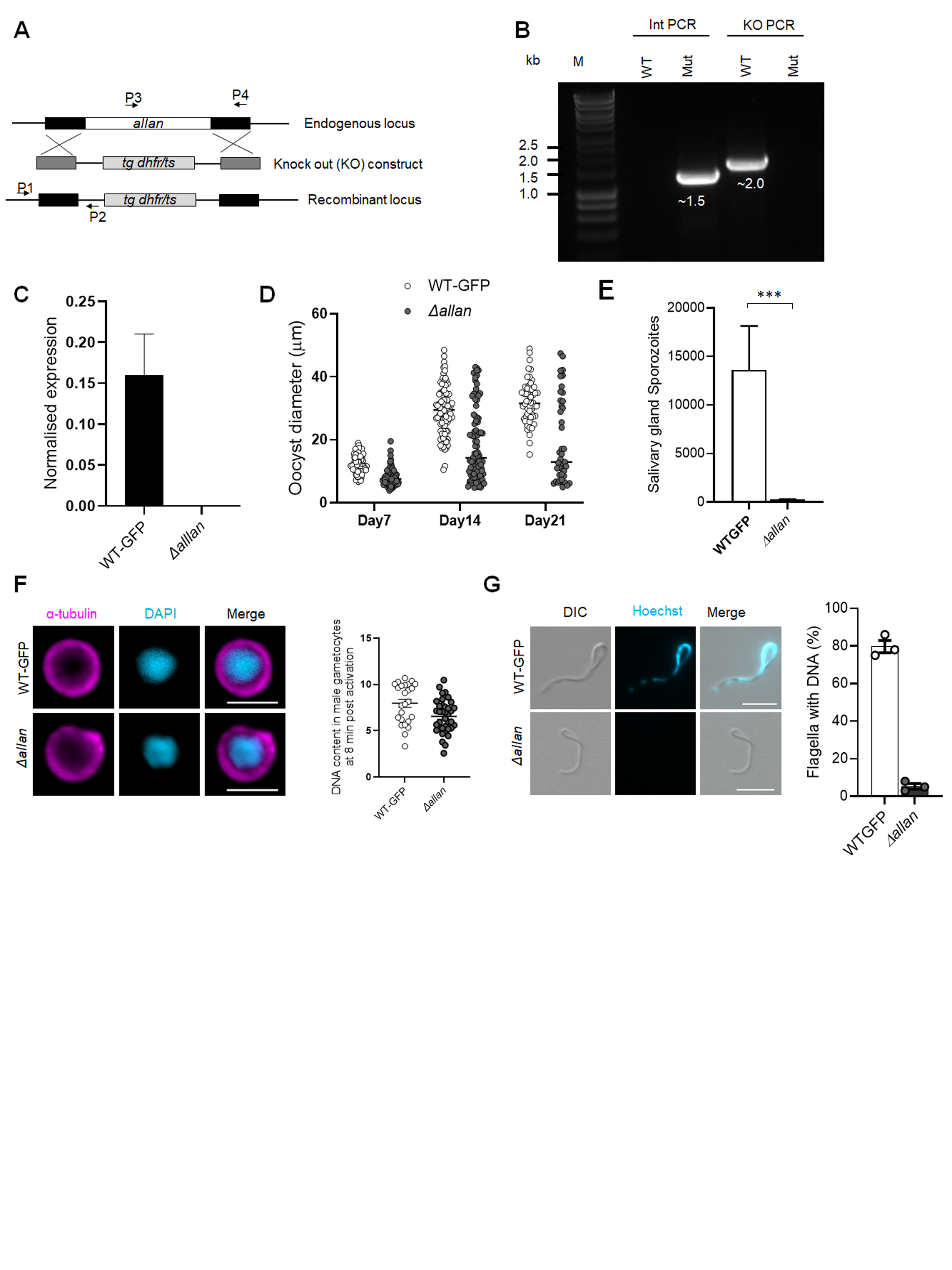

### Fig S11

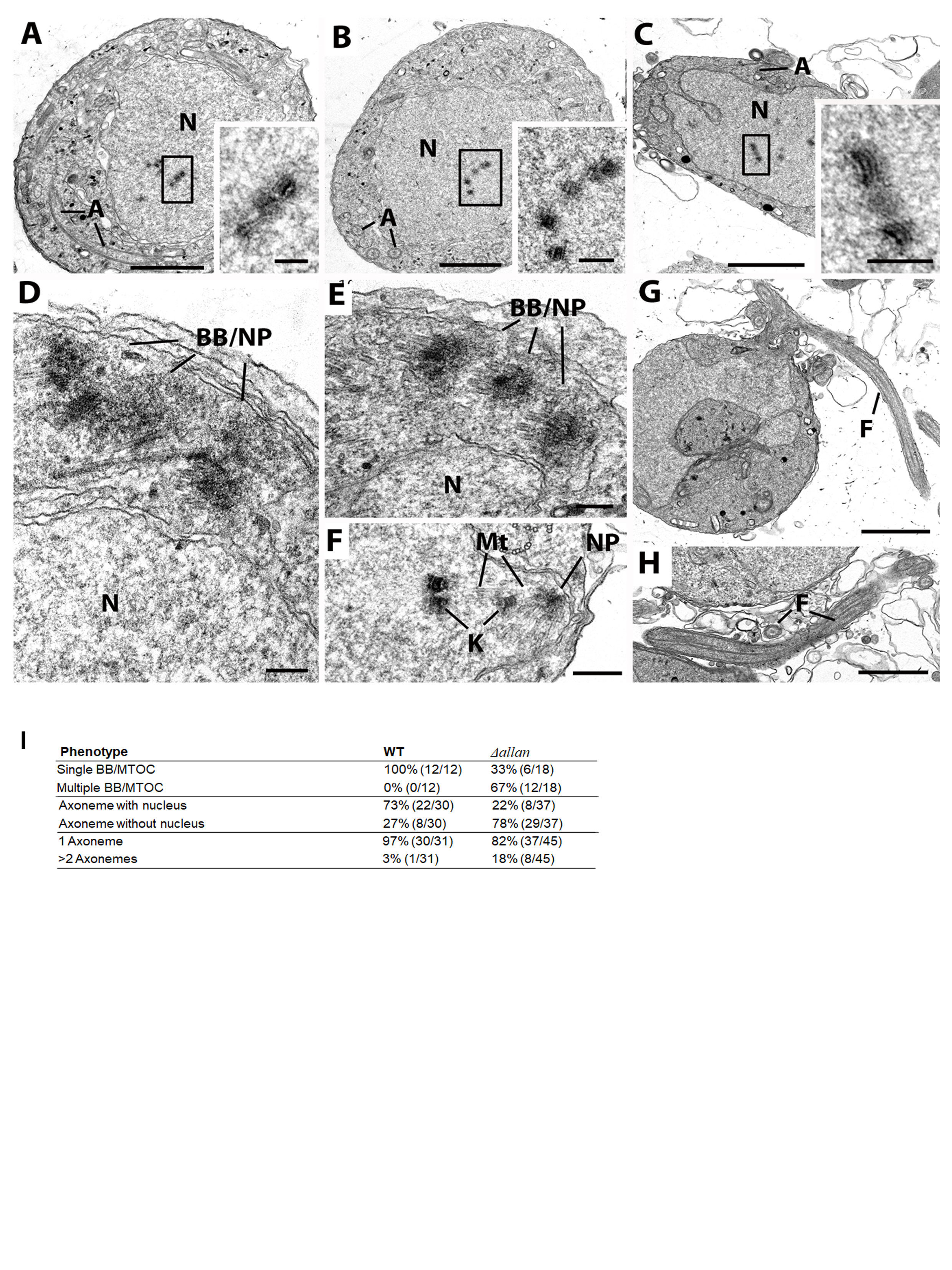
